# Condensin II activation by M18BP1

**DOI:** 10.1101/2024.05.02.592151

**Authors:** Alessandro Borsellini, Duccio Conti, Erin Cutts, Rebecca J. Harris, Kai Walstein, Andrea Graziadei, Valentina Cecatiello, Tom F. Aarts, Ren Xie, Abdelghani Mazouzi, Sushweta Sen, Claire Hoencamp, Richard Pleuger, Sabrina Ghetti, Lina Oberste-Lehn, Dongqing Pan, Tanja Bange, Judith H.I. Haarhuis, Anastassis Perrakis, Thijn R. Brummelkamp, Benjamin D. Rowland, Andrea Musacchio, Alessandro Vannini

## Abstract

Condensin complexes promote the drastic spatial rearrangement of the genome upon mitotic entry. Condensin II initiates chromosome condensation in early mitosis. To prevent chromosome condensation during interphase, condensin II is inhibited by MCPH1, but the mechanism is unknown. Through genetic and proteomic approaches, we identify M18BP1, a protein previously associated with centromere identity, as a factor required for condensin II localization to chromatin. M18BP1 directly binds condensin II’s CAP-G2 subunit and competes with MCPH1 for binding. Upon mitotic entry, CDK1 mediated phosphorylation may promote a switch from MCPH1 to M18BP1 binding to activate condensin II. Our results identify a fundamental and evolutionarily conserved mechanism of condensin II activation.

## INTRODUCTION

During mitosis, the eukaryotic genome must be compacted, spatially organised, and evenly dispatched to two daughter cells. Condensin complexes play a pivotal role in the compaction of chromosomes during mitosis. Two distinct condensin complexes exist in metazoans, condensin I and II. These complexes share their coiled-coil SMC subunits, SMC2 and SMC4, but utilise a different set of kleisin and heat-repeat containing subunits. Condensin I contains CAP-H, CAP-D2 and CAP-G, whereas condensin II contains CAP-H2, CAP-D3 and CAP- G2^1^.

Condensin I and II exhibit distinct subcellular localization and dynamics. During interphase, condensin II is nuclear while condensin I is mostly cytosolic and gains access to chromatin only after nuclear envelope breakdown^2–5^. Condensin II initiates chromosome condensation by shortening chromosomes, after which condensin I reduces chromosome width ^6–9^. Condensin II is also enriched at kinetochores ^3,10–14^, promoting chromatin assembly in the underlying centromeric regions, in turn facilitating kinetochore-microtubule attachment and errorless chromosome segregation ^15–20^. As condensin II is nuclear throughout the cell cycle, it must be kept in check to limit its activation to mitotic entry. Key to this regulation is MCPH1, which prevents condensin II from stably binding to chromatin during interphase by an unknown mechanism. Deletion of the *MCPH1* gene results in condensin II activation and chromosome condensation during interphase^21^.

Here, using orthogonal approaches, we identify MIS18-binding protein 1 (M18BP1) as a determinant of condensin II localization to chromatin. During the G1 phase of the cell cycle, M18BP1 acts as part of a specialized hetero-octameric complex comprising MIS18α and MIS18Β (the MIS18 complex) which associates with the histone chaperone HJURP and the kinase PLK1^22,23^. Together, these proteins ensure that the histone H3 variant CENP-A, an epigenetic marker of centromere specification, is newly deposited in early G1 to compensate for its 2-fold dilution during DNA replication ^10,18,24–31^. Our data reveal that M18BP1 has an additional role, which is crucial for the activity of condensin II. M18BP1 binds directly to the condensin II subunit CAP-G2 and competes with MCPH1 for binding. Therefore, we propose that the role of MCPH1 is to counteract M18BP1, thereby inhibiting condensin II and maintaining the interphase genome in its uncompacted state. Upon mitotic entry, CDK1 activity mediates a switch in binding from MCPH1 to M18BP1, thus triggering condensin II localization to chromatin and ensuring timely chromosome condensation.

## RESULTS

### M18BP1 interacts with condensin II

To identify new regulators of condensin II, we performed a synthetic lethality screen in haploid HAP1 cells already deficient for condensin I (ΔCAP-H), reasoning that they will likely be dependent on condensin II and its regulators (Figure 1A-B). This screen revealed that M18BP1, while not strictly essential in wild type cells, is instead required for the fitness of cells lacking condensin I (Figure 1C).

**Figure 1:**
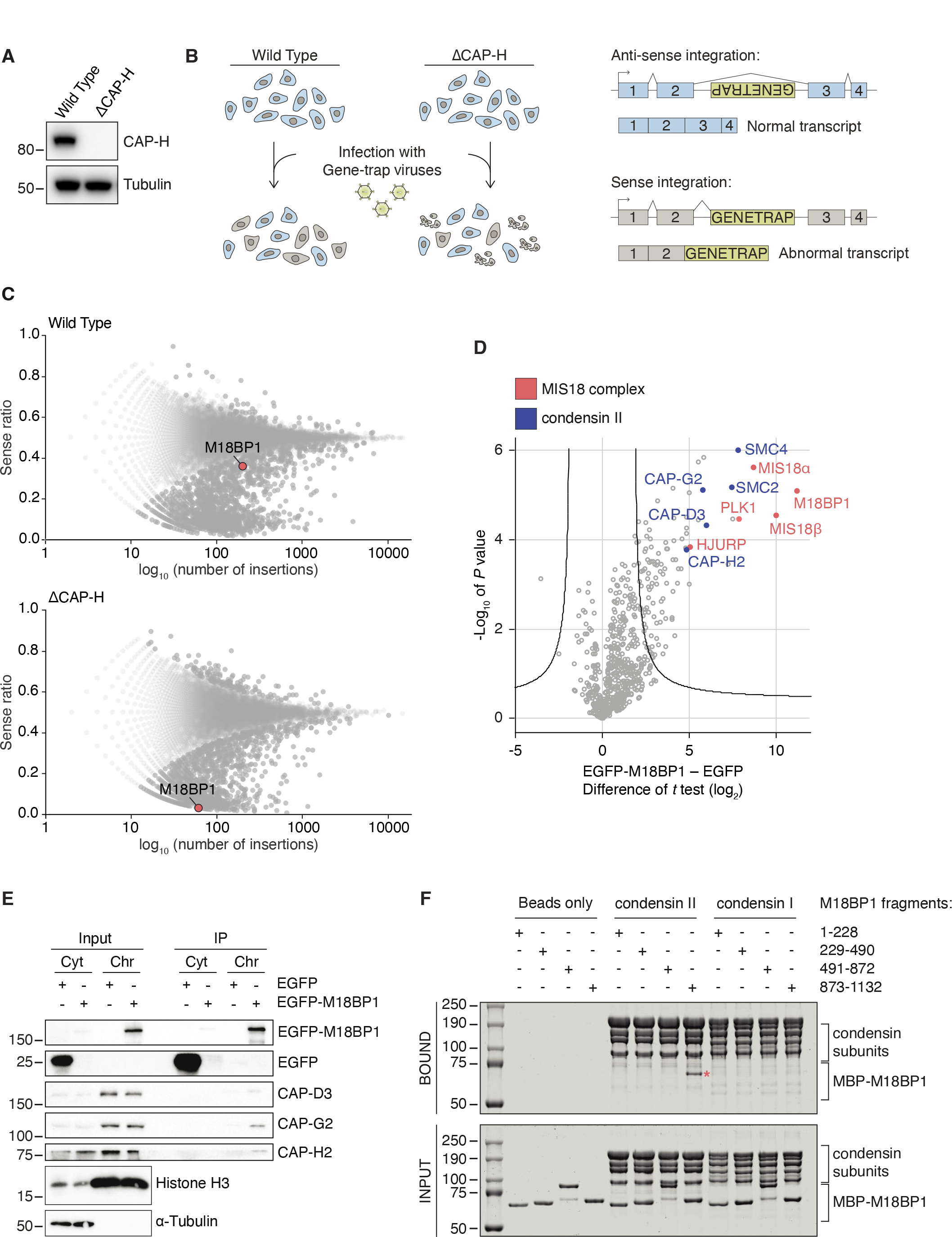
M18BP1 is a condensin II binding partner. (A) Western blot validating that CAP-H is knocked out in the indicated cell line. (B) Schematic of the haploid genetic screening method used in C. (C) Results of the haploid genetic screen. Each dot represents a gene, the x axis shows the number of genetrap virus insertions and the y axis shows the ratio of sense and antisense insertions within individual genes. A ratio of ∼0.5 indicates a non-essential gene. Lower values indicate that the gene is essential for fitness within that genetic background. (D) Volcano plot showing the chromatin interactome of M18BP1 in early G1 phase. Condensin II subunits are marked in blue whereas the CENP-A deposition machinery components are marked in red. (E) Chromatin-bound EGFP-M18BP1 co-immunoprecipitates condensin II in mitotic HeLa cells. (F) *In vitro* pull-down using indicated fragments of MBP-tagged M18BP1 with human condensin I or II. Asterisk indicates the pulled-down M18BP1 fragment.

In parallel, we performed immunoprecipitation (IP) of an EGFP-M18BP1 bait in chromatin extracts from HeLa cells and analysed bound proteins by mass spectrometry (MS) ^29^. As expected, we identified all components of the MIS18 complex^24,27–30,32–35^. In addition, we identified all the subunits of condensin II, but not of condensin I, as highly significant interactors of M18BP1 (Figure 1D). This is consistent with our earlier MS-based identification of M18BP1 in precipitates of an antibody against CAP-H2^36^. An essentially identical list of binding partners was identified using an mCherry-MIS8α bait (Figure S1.1).

To verify the interaction with an orthogonal strategy, we performed a co-IP using EGFP-tagged M18BP1 and confirmed by Western blotting that all components of the condensin II complex are pulled down by M18BP1 from mitotic cell lysates (Figure 1E). To assess whether M18BP1 and condensin interact directly, we performed an *in vitro* pull-down experiment with four MBP-tagged recombinant fragments of M18BP1 and either condensin I or condensin II (Figure 1F). The M18BP1 fragment encompassing residues 873-1132 (M18BP1_873-1132_) was efficiently pulled down by recombinant condensin II, but not by condensin I (Figure 1F). Thus, M18BP1_873-1132_ binds condensin II directly, identifying M18BP1 as a novel condensin II interacting protein.

### M18BP1 uses a short linear motif to engage CAP-G2

We next sought to map the binding site for M18BP1 on condensin II. In contrast to the condensin II holocomplex, condensin II lacking the CAP-G2 subunit was unable to interact with M18BP1_873-1132_ (Figure 2A). In addition, a subcomplex consisting of only CAP-G2 and CAP-H2 was sufficient to pull down M18BP1_873-1132_ (Figure 2A). Together this indicates that the CAP-G2 subunit is key to the M18BP1 interaction.

**Figure 2:**
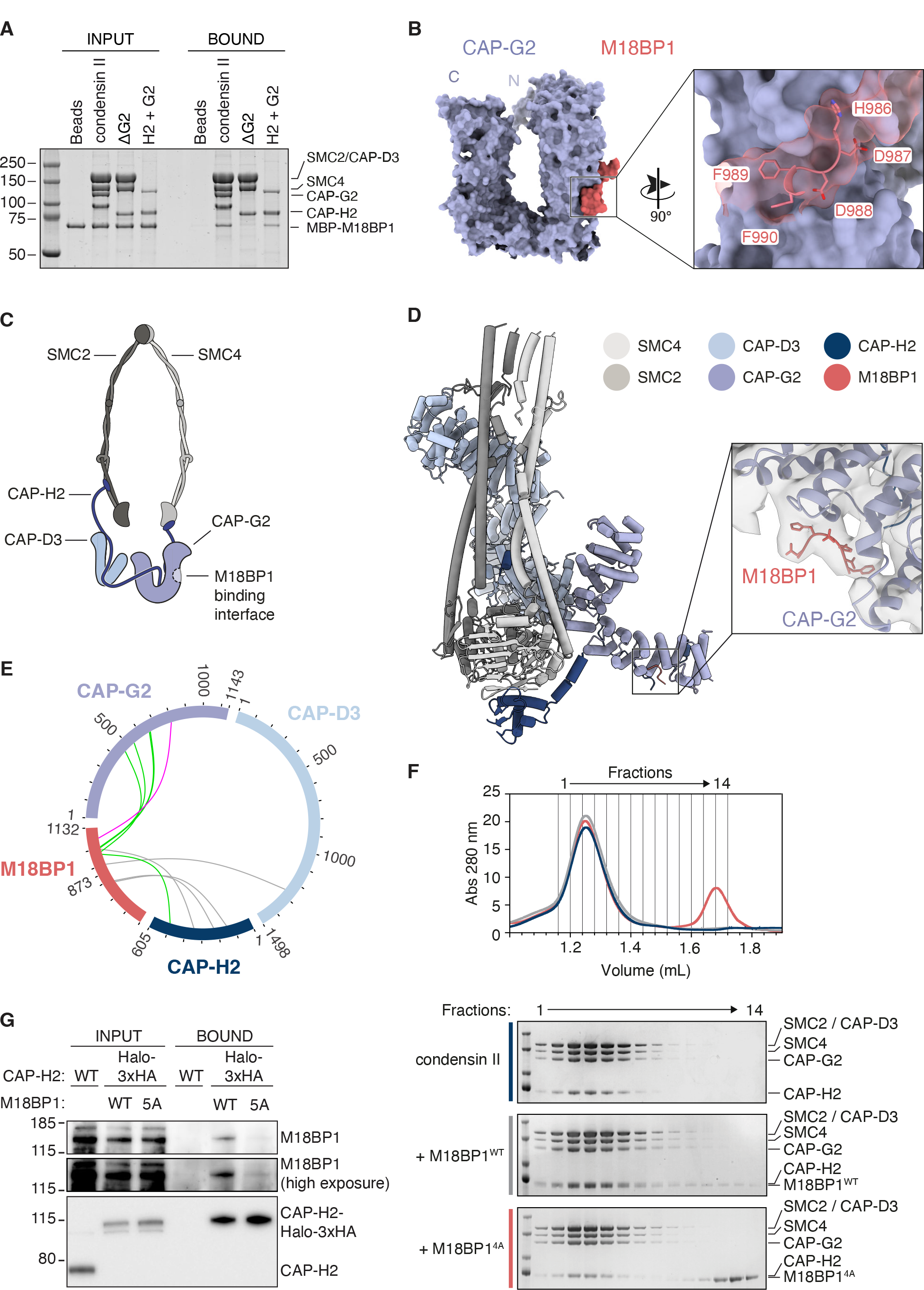
M18BP1 binds to the condensin II CAP-G2 subunit. (A) Pull-down experiment on condensin II strep tag in presence of MBP-M18BP1_873-1132_. In the experiment were used the full length condensin II complex, the complex missing CAP-G2 (ΔG2) or CAP-H2 and CAP- G2 subunits only. (B) AF2 multimer prediction of CAP-G2 (purple) with M18BP1 (red), with a zoom-in of the “HDDFF” motif of M18BP1. (C) Cartoon representation of the condensin II complex. (D) Cryo-EM structure of apo condensin II holo-complex displaying SMC subunits in grey, CAP-D3 in light blue, CAP-G2 in purple and CAP-H2 in dark blue and a zoom-in of the extra density at the CAP-G2 - M18BP1 (red) interaction interface. (E) Inter-molecular sulfo-SDA crosslinks between M18BP1_873-1132_ and condensin II subunits. (F) Size exclusion profiles of condensin II (blue), condensin II with M18BP1_WT-873-1132_ (grey) and condensin II with M18BP1_4A-873-1132_ (red). Fractions from size exclusion profiles are shown in the SDS page gels in the lower panels. (G) Western blot analysis after co-immunoprecipitation using anti- HA beads in wild type cells expressing endogenous untagged CAP-H2 (control cells), wild type cells expressing endogenously tagged CAP-H2-Halo-3xHA or M18BP1_5A_ (“HDDFF” mutant) cells expressing CAP-H2-Halo-3xHA.

To further characterize the binding interface between M18BP1 and condensin II, we identified highly conserved patches within the M18BP1_873-1132_ fragment and mutated blocks of five residues (Figure S2.1A) to alanine. This identified M18BP1 residues 984-988 (corresponding to the motif Asp-Asp-His-Asp-Asp, or DDHDD) as being necessary for binding (Figure S2.1B). In line with this, an AlphaFold-Multimer (AF2) ^37,38^ structural model predicts with high confidence a binding interface between CAP-G2 and M18BP1 in which a partly overlapping linear motif of M18BP1 comprising residues 986-990 (His-Asp-Asp-Phe-Phe, or HDDFF) (Figure S2.1C) makes extensive contacts with CAP-G2 (Figure 2B-C and Figure S2.1D-E).

To visualize the molecular basis of the interaction between M18BP1 and condensin II in further detail, we determined the cryo-EM structure of condensin II with M18BP1_873-1132_ at 7 Å average resolution, with a local resolution ranging between 5 and 11 Å. Individual subunits of the holo-complex, generated by AF2, could be confidently fitted to the EM maps (Figure 2D and Figures S2.2-S2.3). In absence of both DNA and ATP, the overall architecture of human condensin II is reminiscent of *S. cerevisiae* condensin in its apo form^39^. The two SMC subunits form a parallel coiled coil pair with the ATPase heads held in close proximity, albeit not in an engaged ATP hydrolysis-competent conformation, and with flexible “elbow” and “hinge” domains of the SMC subunits, which fade out of density.

Although most of M18BP1 is not visible in the EM map, an unassigned density could be observed exactly at the position where AF2 predicts the interaction interface between the M18BP1 “HDDFF” motif and the CAP-G2 subunit (Figure 2D). Crosslinking coupled with mass-spectrometry (crosslinking MS) of the condensin II-M18BP1_873-1132_ complex supported the overall architecture of the complex obtained by cryo-EM, with intermolecular crosslinks consistent with the conformation adopted by the heat repeats in our structure and with the interaction with the CAP-H2 kleisin subunit (Figure S2.4). Importantly, the crosslinking data also support the interaction interface between M18BP1 and CAP-G2 (Figure 2E), with a specific crosslink between CAP-G2 K496 and M18BP1 L995, close to the 986-HDDFF-990 motif. Thus, the “HDDFF” motif of M18BP1 is implicated in condensin II binding. We therefore mutated this motif to “HAAAA” in the M18BP1_873-1132_ fragment (M18BP1_4A_) and monitored co-elution with condensin II in size-exclusion chromatography. While the wild type M18BP1_873-1132_ fragment co-eluted with condensin II, the same fragment harbouring the “HAAAA” mutation failed to bind and eluted in a separate peak (Figure 2F).

Next, by modifying the endogenous *M18BP1* locus using CRISPR-Cas9 technology in HAP1 cells, we generated an M18BP1 mutant with all five residues in the “HDDFF” motif modified to alanine (M18BP1_5A_). In addition, we endogenously tagged the condensin II subunit CAP-H2 with a Halo-3xHA tag in both wild type and M18BP1_5A_ cells (Figure S2.5). Immunoprecipitated CAP-H2-Halo-3xHA bait from these cells pulled down wild type M18BP1 but not M18BP1_5A_ (Figure 2G). Taken together, these results indicate that M18BP1 and condensin II form a stable complex that is mediated, partly or entirely, by an interaction between a conserved M18BP1 linear motif and the condensin II subunit CAP-G2.

### M18BP1’s “HDDFF” motif localizes condensin II to chromatin

To investigate the role of M18BP1’s binding to condensin II, we used the endogenously mutated M18BP1_5A_ cells which also express CAP-H2 tagged with Halo-3xHA. As expected, in wild type cells CAP-H2 was highly enriched on chromatin during mitosis. Remarkably, in M18BP1_5A_ cells CAP-H2 was almost undetectable on chromosomes of prometaphase-arrested cells (Figure 3A-B). This observation was corroborated by imaging untagged CAP-D3 in HeLa cells expressing siRNA-resistant, mCherry-tagged wild type M18BP1 or M18BP1_4A_ mutant. After arrest in metaphase using MG132, CAP-D3 was highly enriched on chromatin in control cells.

**Figure 3:**
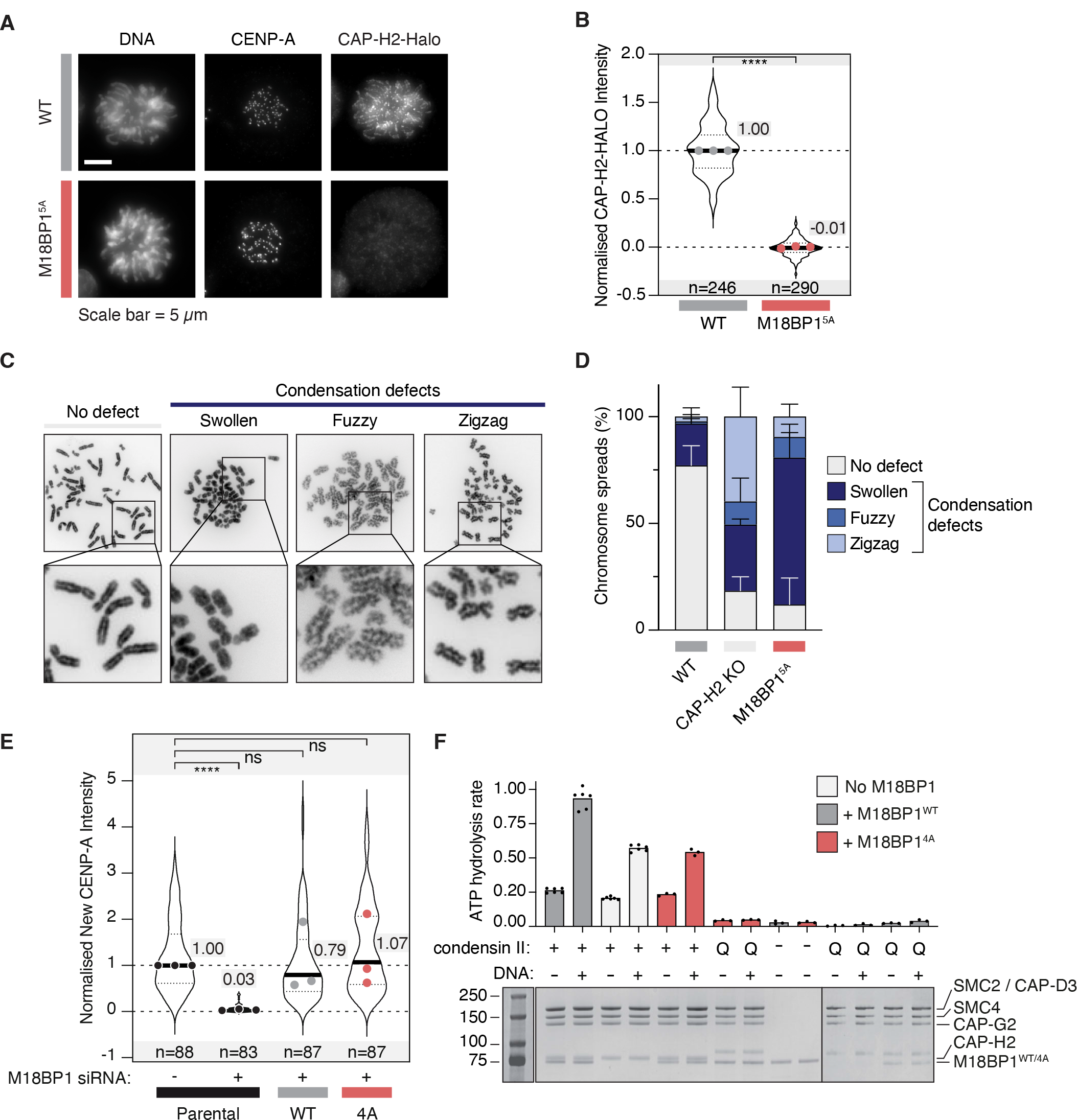
M18BP1 is a major recruiter of condensin II during mitosis. (A) Representative images of endogenously tagged CAP-H2-Halo-3xHA levels on mitotic chromatin in wild type (WT) or M18BP1_5A_ “HDDFF” mutant cells. (B) Quantification of CAP-H2-HALO levels from A. Dots show the median of each experimental repeat. “n” represents the total number of cells from three independent experiments. The median of the combined data is shown for each condition and dots show the median of each experimental repeat. (C) Example images of mitotic chromosome spreads with condensation defects as quantified in D. (D) The percentage of chromosome spreads with condensation defects in wild type, ΔCAP-H2 and M18BP1 HDDFF mutant cells. Bars represent mean ± SD. (E) The M18BP1_4A_ mutant does not affect new CENP-A deposition. Plot shows the intensity of the newly deposited CENP-A in the indicated conditions. “n” represents the total number of cells from three independent experiments. The median of the combined data is shown for each condition and dots show the median of each experimental repeat. (F) ATPase rate of the condensin II complex in presence of M18BP1_873-1132_ and DNA. Q refers to Condensin II with an ATPase deficient mutation in the Q-loop. 4A refers to the M18BP1_4A_ mutant. Below is an SDS page gel showing the loading controls. Black dots are individual values from 3 to 6 independent experiments.

Conversely, CAP-D3 was no longer detected on chromosomes depleted of M18BP1 by RNAi (Figure S3.1A-C). Ectopic expression of wild type M18BP1 rescued CAP-D3 localization to mitotic chromosomes, whereas expression of M18BP1_4A_ failed to do so (Figure S3.1A-B, D-E). The depletion of M18BP1 prior to mitotic entry did not affect the overall levels or nuclear localization of CAP-D3 in G2 cells (Figure S3.2). Together, these results show that M18BP1 plays a central role in condensin II association with chromatin during mitosis, and that this role requires the M18BP1 “HDDFF” motif.

As M18BP1 mutant cells have no detectable condensin II on mitotic chromosomes (Figure 3A and Figure S3.1), we used chromosome spreads to determine whether these cells experienced condensation defects. As expected, wild type cells showed well-compacted mitotic chromosomes, whereas cells lacking functional condensin II (ΔCAP-H2 cells) showed poorly condensed chromosomes. M18BP1_5A_ cells displayed condensation defects to a similar extent as ΔCAP-H2 cells (Figure 3C-D), although the specific morphology of the chromosomes somewhat differs, for reasons that remain unclear. Consistent with these results, inter-kinetochore distances on congressed chromosomes experiencing microtubule-generated forces were significantly higher in HeLa cells depleted of M18BP1 relative to non-depleted controls (Figure S3.3). Removal of the condensin II subunit CAP-G2 using siRNAs also resulted in a similar increase (Figure S3.3). Together, these results corroborate the idea that M18BP1 is required for condensin II-mediated chromosome condensation.

Since M18BP1 is essential for the deposition of the centromeric marker CENP-A ^24,29,31,40^, we asked whether disruption of the “HDDFF” motif also affects this pathway. Specific labelling of newly deposited CENP-A showed that M18BP1_4A_ is as efficient in depositing CENP-A as wild type M18BP1 (Figure 3E and Figure S3.4). Thus, mutating the M18BP1 “HDDFF” motif yields a separation of function mutant that ablates condensin II localization to mitotic chromosomes but does not affect the process of CENP-A deposition.

### M18BP1 enhances condensin II DNA-dependent ATPase activity

Condensin II-mediated chromosome condensation is dependent on its ATPase activity ^41,42^. We therefore asked whether M18BP1 might affect condensin II ATPase activity. In line with previous evidence^43^, DNA stimulated the ATP hydrolysis rate of condensin II by ∼2.5 fold and addition of M18BP1_873-1132_ further stimulated the ATP hydrolysis rate, but the effect was less pronounced if DNA was omitted (Figure 3F). The M18BP1_4A_ mutant did not enhance condensin II ATPase activity, however, indicating that a direct interaction between M18BP1 and condensin II is required to stimulate ATP hydrolysis. A condensin II Q-loop mutant (SMC2 Q147L and SMC4 Q229L), which is deficient in ATP hydrolysis ^44–49^, did not show unspecific ATPase activity, even upon addition of DNA or of M18BP1 (Figure 3F).

### M18BP1 competes with the condensin II antagonist MCPH1

MCPH1 binds the CAP-G2 subunit of condensin II using a conserved “central domain” ^21^. We noted that a highly conserved “YDDYF” motif within the MCPH1 central domain is remarkably similar to the “HDDFF” motif of M18BP1 (Figure 4A), thus suggesting that MCPH1 and M18BP1 may bind the same interface of condensin II as competitive agonists. Confirming this idea, AF2 predicted with high confidence that the “YDDYF” and “HDDFF” motifs of MCPH1 and M18BP1, respectively, bind the same interface of CAP-G2 (Figure 4B and Figure S4.1). Crosslinking MS of condensin II in complex with MCPH1_1-435_ identified multiple crosslinks connecting the MCPH1 central domain to the predicted interface of CAP- G2, thus validating the AF2 prediction (Figure 4C). We then used fluorescence polarisation competition assays to assess the prediction that MCPH1 and M18BP1 compete for condensin II binding. Using a MCPH1_407-424_ 5-FAM-labelled peptide and condensin II at a fixed concentration and increasing concentrations of unlabelled M18BP1 or MCPH1, we confirmed that M18BP1 and MCPH1 compete for condensin II binding (Figure 4D).

**Figure 4:**
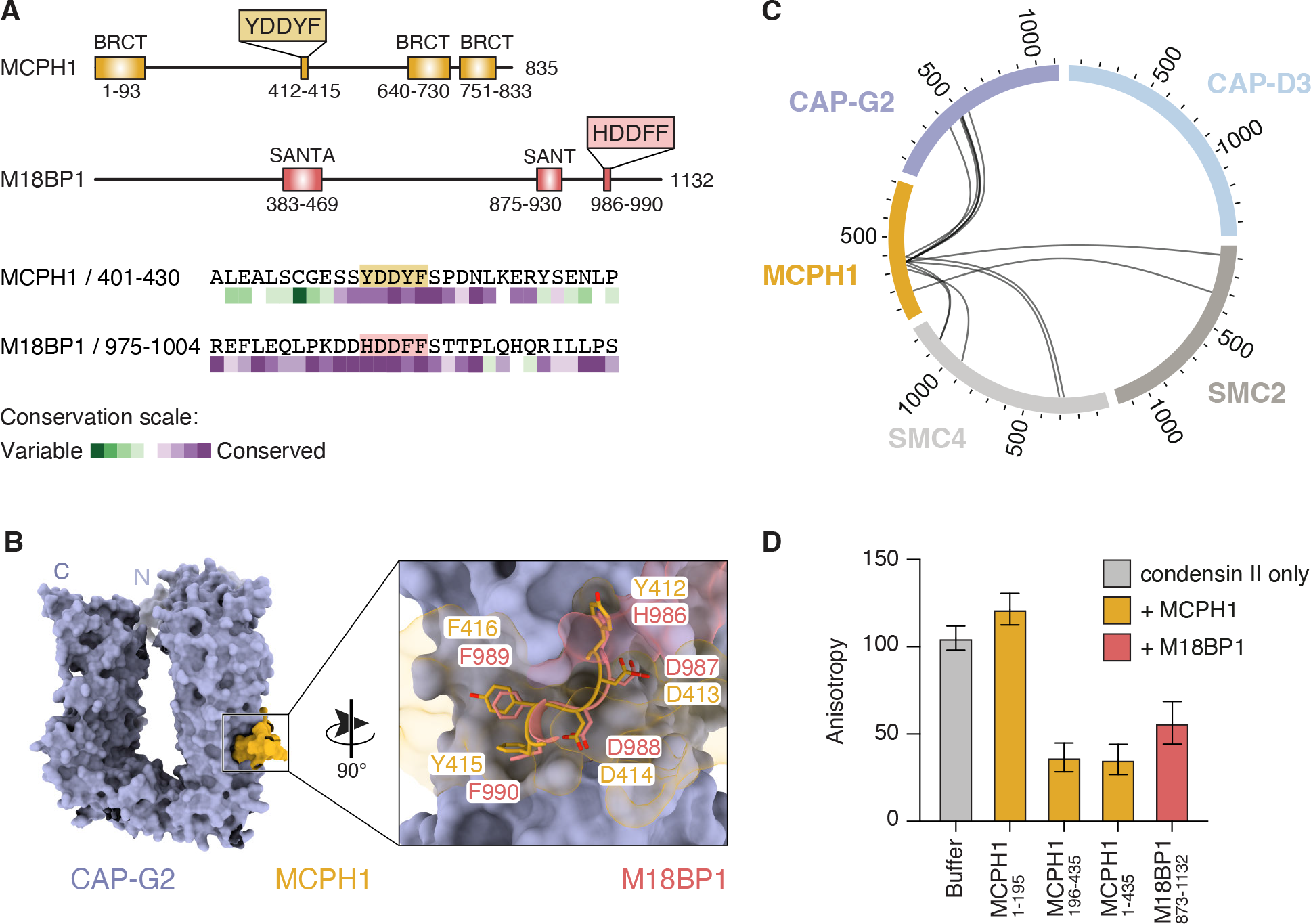
M18BP1 and MCPH1 bind to the same site on CAP-G2. (A) Schematic of M18BP1 and MCPH1 indicating the identified binding motifs. Below shows the alignment of the motifs found in M18BP1 and MCPH1, colored by conservation score from ConSurf. (B) AF2 Multimer structural prediction of CAP-G2 bound to MCPH1 and M18BP1. The two predictions were overlaid using CAP-G2 as the reference. For clarity, only the YDDYF/HDDFF motifs are shown. (C) Crosslinking mass spectrometry of MCPH1 bound to condensin II. (D) Fluorescence anisotropy competition assay using a 5-FAM-MCPH1_407-424_ peptide probe at 0.3 µM and a fixed concentration of condensin II at 0.8uM, with indicated MCPH1 or M18BP1 fragments added to compete with the probe. Mean and standard error from n=3 repeats.

### M18BP1 is essential for interphase condensation in absence of MCPH1

Although our biochemical observations identify M18BP1 and MCPH1 as competitive agonists of condensin II *in vitro*, whether competition occurs *in vivo* is unclear. To address this question, we first assessed if these proteins are differentially regulated during the cell cycle. In both HeLa and RPE1 cells, M18BP1 was strongly enriched at centromeres during early G1 phase, as observed previously^24^ (Figure 5A-B and Figure S5.1). Additionally, we also observed substantial levels of M18BP1 in S-phase and G2 at both the centromeres and diffusely throughout the nucleus (Figure 5A-B, S5.1). Thus, M18BP1 localizes to chromatin well before mitotic onset, suggesting that it could compete with MCPH1 for condensin II binding. Nonetheless, condensin II does not associate stably with chromatin until the onset of mitosis, likely because the repressive activity of MCPH1 keeps it in check during interphase^21^. In cells lacking MCPH1, condensin II stably binds to chromatin during interphase, resulting in interphase chromosome condensation^21^.

**Figure 5:**
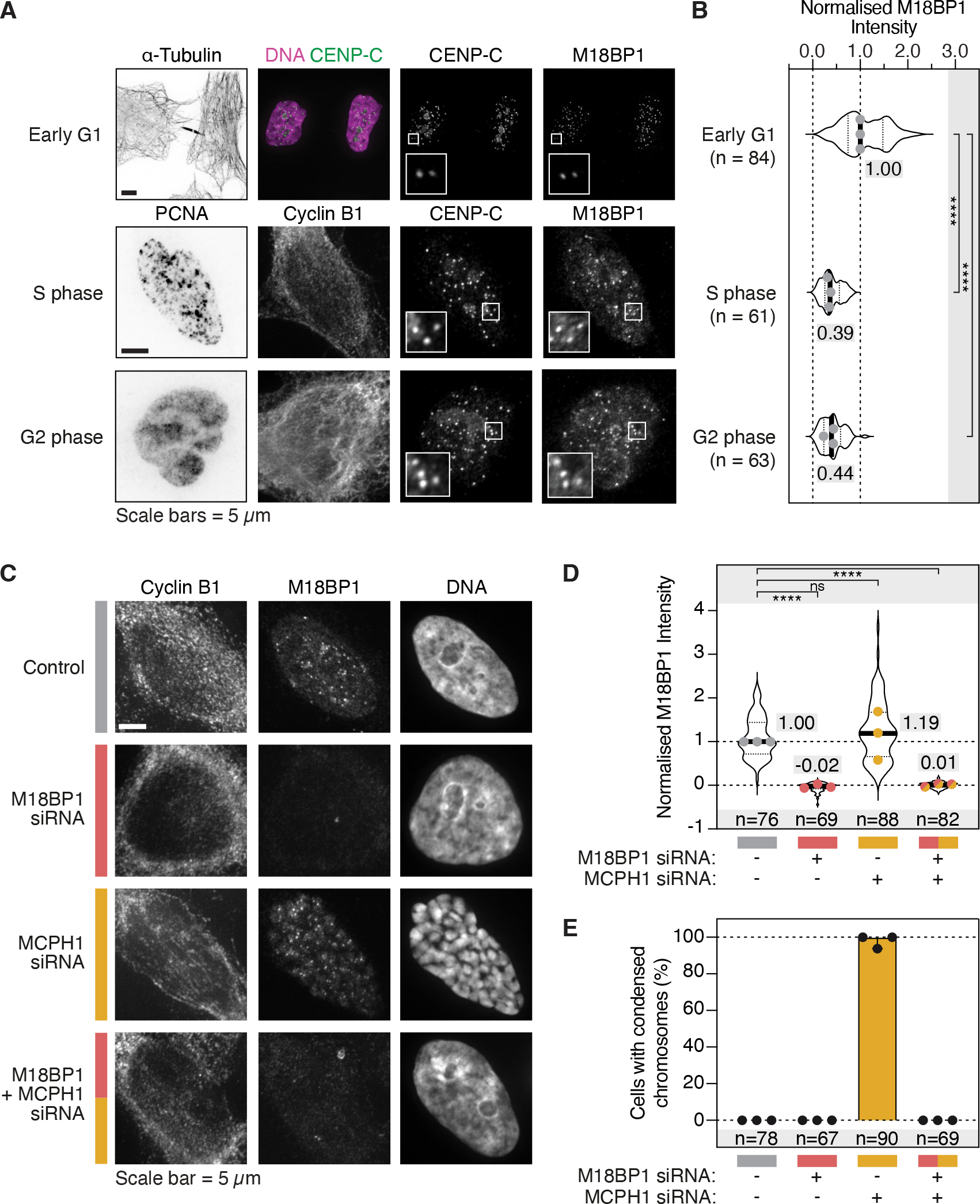
M18BP1 is essential for interphase condensation in the absence of MCPH1. (A) Localisation of M18BP1 on chromatin and at centromeres during the cell cycle in HeLa cells. Endogenous M18BP1 was detected using an antibody. Different cell cycle stages were identified using α-Tubulin, PCNA and CyclinB1 signals. Insets show a zoom in on the centromeres. (B) Quantification of centromeric M18BP1 levels from A. “n” represents the number of cells analysed across three independent experimental repeats. The median of the combined data is shown for each condition and dots show the median of each experimental repeat. (C) Example images of chromosome condensation status of HeLa cells in G2 when treated with different combinations of siRNA. (D) Quantification of M18BP1 chromatin levels from C. “n” represents the number of cells analysed across three independent experimental repeats. The median of the combined data is shown for each condition and dots show the median of each experimental repeat. (E) Quantification of the chromosome condensation status of cells from C. Bars and whiskers indicate the median and 95% confidence interval, respectively.

At least two mutually exclusive scenarios may underlie the interphase condensation observed in absence of MCPH1. In the first scenario, condensin II is sufficient for autonomous localization to chromatin, and M18BP1 is merely required to relieve repression by MCPH1 as cells enter mitosis. In the second scenario, condensin II requires M18BP1 to localize to chromatin, and MCPH1 prevents M18BP1 from acting during interphase, limiting condensin II activity to mitosis. To distinguish between these two possibilities, we utilised the interphase condensation phenotype observed in MCPH1 deficient cells ^21^. As expected, chromosome condensation was observed in G2 cells upon depletion of MCPH1 (Figure 5C-D and Figure S5.2). Importantly, condensation was completely suppressed when M18BP1 was also depleted (Figure 5C-E). Thus, M18BP1 is required for condensin II mediated chromosome condensation in interphase cells lacking MCPH1. Likewise, M18BP1 is required for chromosome condensation in MCPH1-depleted mitotic cells, as condensin II recruitment was lost once M18BP1 was co-depleted (Figure S5.3). This suggests that M18BP1 does not merely interact with condensin II to counteract MCPH1. Instead, our data support a scenario in which M18BP1 plays a central role in condensin II’s ability to condense chromatin, and in which MCPH1 prevents interphase condensation by counteracting M18BP1.

### Phosphorylation may trigger a switch from MCPH1 to M18BP1

A closer look at the condensin II binding motifs in MCPH1 and M18BP1 revealed that both motifs display adjacent canonical S/TP consensus sites for the mitotic kinase CDK1 (Figure 6A). AF2 structural models of the CAP-G2-M18BP1 and CAP-G2-MCPH1 complexes suggest how CDK1 driven phosphorylation might impact their relative binding affinity for condensin II (Figure 6B). MCPH1 S417 is predicted to establish multiple contacts with other MCPH1 residues required to stabilize a kinked helix involved in CAP-G2 binding. Phosphorylation of this residue likely destabilises the interaction with CAP-G2. In agreement with this model, previous work has demonstrated that phosphorylation of MCPH1 S417 leads to a reduced affinity for condensin II^21^. By contrast, the putative CDK1 phosphorylation site of M18BP1, T993, neighbours a positively charged region of CAP-G2. Phosphorylation of this residue may enable an additional electrostatic interaction, increasing the binding affinity for condensin II.

**Figure 6:**
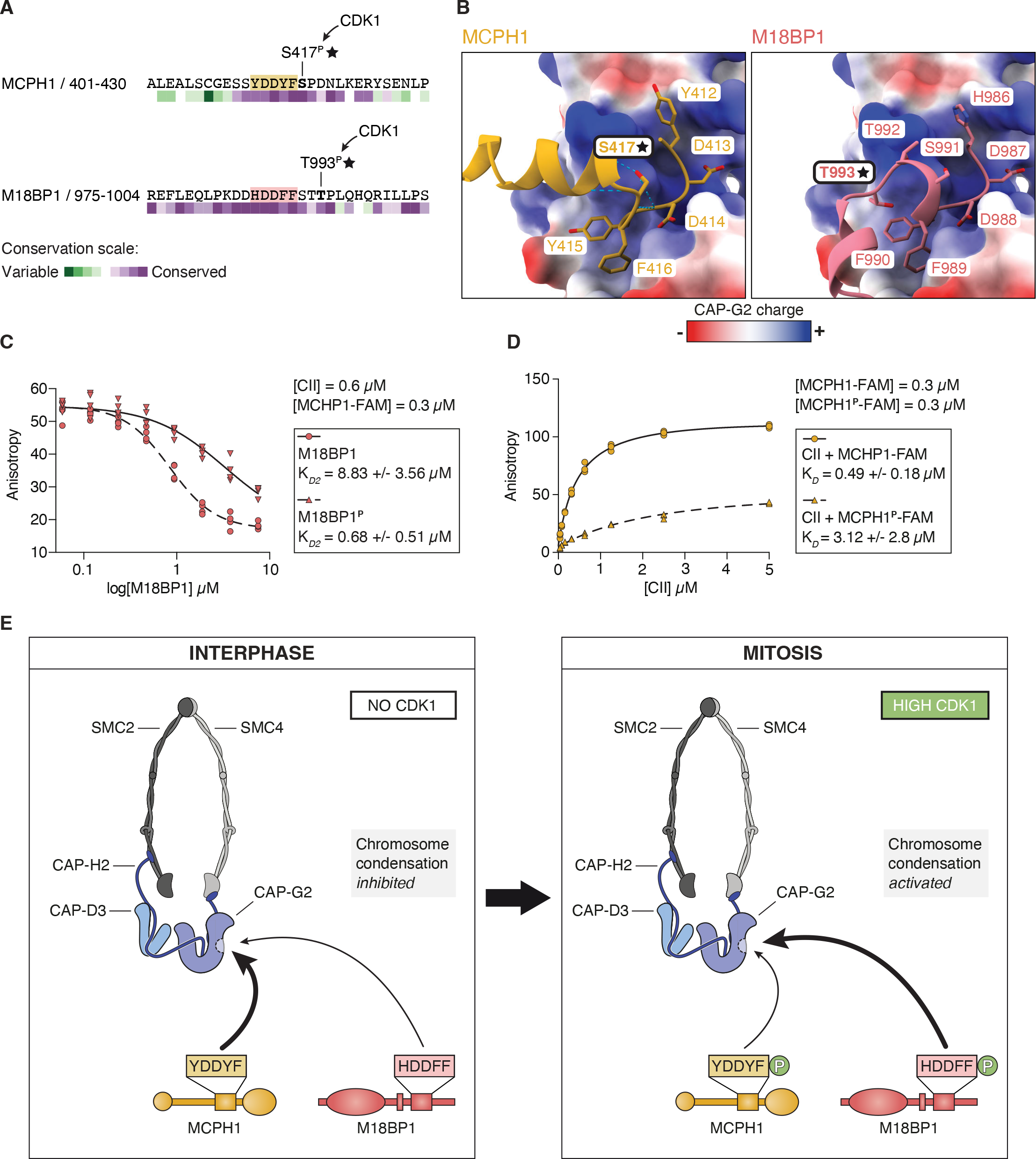
A phosphorylation-driven switch from MCPH1 to M18BP1. (A) Predicted CDK1/cyclin B kinase phosphorylation sites adjacent to MCPH1-YDDYF and M18BP1- HDDFF motifs. (B) Close-ups of AlphaFold2 predictions of MCPH1 (yellow) and M18BP1 (red), respectively, with CAP-G2 which is colored by electrostatic potential. Black stars indicate the location of the potential phosphorylation sites. Blue dotted lines indicate predicted H bonds. (C) Fluorescence anisotropy binding assay with increasing concentrations of condensin II and a fixed concentration (0.3 µM) of wild-type or phosphorylated MCPH1_407-424_ peptides. (D) Fluorescence anisotropy competition assay using a 5-FAM-MCPH1_407-424_ peptide and condensin II at a fixed concentration (0.3 µM and 0.6 µM, respectively), with increasing concentration of wild type or phosphorylated M18BP1_873-1132_. M18BP1 was phosphorylated by incubation with CDK1-Cyclin B. (E) Model of condensin II regulation by MCPH1 and M18BP1. During interphase (left panel), MCPH1 binds to CAP-G2 and inhibits condensin II activity. Upon mitotic entry (right panel), high CDK1 activity induces phosphorylation of both MCPH1 and M18BP1. This leads to a switch from CAP-G2 binding MCPH1 to M18BP1 thus activating condensin II and initiating chromosome condensation.

To assess how phosphorylation affects M18BP1 and MCPH1 binding to condensin II, we performed fluorescence polarisation competition assays using a fixed concentration of condensin II and 5-FAM labelled MCPH1 peptide. Pre-incubation of M18BP1_873-1132_ with CDK1 increased the affinity of M18BP1_873-1132_ for condensin II approximately 10-fold (Figure 6C). Importantly, crosslinking MS identifies crosslinks between peptides containing the phosphorylated form of M18BP1 T993 and CAP-G2, confirming that CDK1-dependent phosphorylation of M18BP1 T993 is a determinant of M18BP1-condensin II interaction. Conversely, a 5-FAM labelled MCPH1 peptide phosphorylated on S417, a bona fide substrate of CDK1, caused a 6-fold reduction in binding affinity compared to the unphosphorylated form of the peptide (Figure 6D). Collectively, these data suggest that CDK1 phosphorylation of M18BP1 and MCPH1 at the onset of mitosis switches the binding preference of condensin II from MCPH1 to M18BP1, triggering condensin II localization to chromatin and the onset of chromosome condensation.

## DISCUSSION

Timely activation of condensin complexes underlies the dramatic compaction of chromosomes at the transition from interphase to mitosis, which is essential for accurate chromosome segregation. How this transition is regulated, however, is poorly understood. Here, we identify M18BP1 as a regulator of condensin II required for chromosome condensation.

### A moonlighting role for M18BP1

M18BP1 is well-known for its role in maintaining centromere-specific chromatin through the deposition of new CENP-A in the G1 phase of the cell cycle^10,26^. CENP-A deposition was previously linked to condensin II through its interaction with HJURP, a specialized chaperone required to load CENP-A onto chromosomes ^18^. We surmise that the interaction of condensin II with HJURP is indirect and mediated by M18BP1 through the MIS18 complex, which binds and recruits HJURP to the kinetochore^28,50^. This interpretation is consistent with our identification of condensin II in precipitates of the MIS18 complex subunit MIS18α, onto which HJURP docks. Thus, the interaction of M18BP1 with condensin II appears to be consistent with its interaction with HJURP and the MIS18 complex.

Depletion of condensin II has also been shown to reduce the loading of new CENP-A at centromeres^16,18^. However, we show here that an M18BP1 mutant impaired in condensin II binding retains the ability to load new CENP-A. Thus, the precise role of condensin II in CENP-A loading requires further investigation. The intimate connection of kinetochores with condensin is further substantiated by a role of CENP-I, a protein associated with CENP-A and required for CENP-A loading, in kinetochore accumulation of condensin^11^. In unpublished work, we have identified CENP-I as part of the M18BP1 receptor at kinetochores (KW, DP, and AM, unpublished results). In addition, many other regulators such as transcription factors, histones, various histone modifications, protein phosphatase 2A (PP2A), Shugoshin, and telomeric proteins have been proposed to contribute to chromosome recruitment of condensin II ^13,51–61^. Whether these determinants are linked to the function of M18BP1 is currently unclear and grants future analyses.

### A molecular switch that triggers chromosome condensation

Collectively, our data point to a model in which M18BP1 is required for condensin II to localize to chromatin and condense chromosomes. During interphase MCPH1 counteracts this activity of M18BP1, directly competing for binding to CAP-G2 and preventing the stable association of condensin II with chromatin, thereby maintaining the interphase genome in its uncompacted state. At the onset of mitosis, CDK1-mediated phosphorylation of both M18BP1 and MCPH1 switches preferences, favouring the binding of M18BP1 over MCPH1, and resulting in condensin II localization to chromatin and chromosome condensation specifically as cells enter mitosis (Figure 6E).

An MCPH1/M18BP1 competition model is compatible with MCPH1 knock-out studies in which the absence of MCPH1 allows for condensin II association with chromosomes during interphase. In this setting, condensin II was highly enriched at centromeres^21^. Both the centromeric enrichment and the overall diffuse chromatin localisation of M18BP1 that we detect in interphase cells, could enable M18BP1 to activate condensin II upon MCPH1 depletion, without the need for CDK1-mediated phosphorylation. How M18BP1 is localized to DNA beyond centromeric regions remains unknown. We speculate that the chromosome localization of M18BP1 in interphase may represent a primed state in which M18BP1 is positioned to allow for explosive condensin II loading in mitosis upon CDK1 activation.

While the existence of a CDK1 controlled switch emerges with clarity from our observations, why the complexes of condensin II with M18BP1 and MCPH1 result in opposite functional outcomes despite closely related binding modes remains unclear. Other regions within M18BP1 and/or MCPH1 may influence condensin II activity in a different manner. Another intriguing observation is that *in vitro* condensin II is capable of DNA binding and compaction in the absence of M18BP1^62^. Why M18BP1 is required for condensin II localization to chromatin in cells, but not *in vitro*, is unclear. Future work will be needed to address these important mechanistic questions.

### Unique structural features of human condensin II

The cryo-EM structure of human condensin II in complex with M18BP1 displays several unique features not observed in the structures of yeast condensin homologues^39,47,63,64^. First, the heat repeats-containing subunits CAP-D3 and CAP-G2 bind simultaneously to the SMC moiety (Figure S6.1A). In particular, while CAP-D3 binds to SMC2 in two positions situated near the ATPase head (residues 104-110) and along the coiled-coil (residues 215-235 and 941- 98), CAP-G2 binds to SMC4 in the vicinity of the W-loop in the ATPase head (residues 1182- 1186). In the structures of yeast condensin, this part of SMC4 is in contact with the conserved KG loop of the CAP-D3 homologue Ycs4^39,47^ (Figure S6.1B), representing a notable difference between the human and yeast complexes. The heat repeats of CAP-G2 and CAP-D3 contact each other near residues 330-342 and residues 1215-1252, respectively, forming almost a continuous heat repeat structure (Figure 2D and S6.1A). The CAP-D3 residues that bind to CAP-G2, named “heat docker” domain (Figure S6.1A), were recently reported to be important for the regulation of condensin II activity in *Xenopus* egg extracts^65^, suggesting that these interactions may play a regulatory role across vertebrates. Deletion of the heat docker resulted in an increased condensin loading phenotype, which led to the hypothesis of a downregulation mechanism in which the CAP-D3 binding to CAP-G2 and/or CAP-H2 locks condensin II into a less active complex^65^.

The kleisin CAP-H2 simultaneously contacts all subunits of the condensin II complex (Figure S6.1C). The CAP-H2-SMC4 interface is comparable to the one observed in the previous structures from yeast homologues^39,47,63,64^. In contrast, the CAP-H2 N-terminal domain, that is flexible in the structure of the apo non-engaged conformation of the *S. cerevisiae* homologue, is sandwiched between the SMC2 coiled-coil neck and one of the CAP- D3 “heat docker” helices. In addition, density for the internal CAP-H2 domains is present on the surfaces of both heat repeats in positions that are consistent with both previous structures of yeast homologues and AF2 predictions (Figures S6.1C).

Superimposition with the structure of DNA-bound yeast condensin^64^ reveals that the position of the subunit CAP-D3 in the apo condensin II cryo-EM structure obstructs accessibility to the DNA path above the ATPase heads. (Figure S6.1D). Thus, it remains to be clarified how ATP binding can rearrange the condensin II complex in a DNA binding-competent conformation. The cryo-EM structure may have captured an intermediate of the loop enlargement cycle between ADP release and ATP binding.

### A universal mechanism to control genome architecture

CDK1 phosphorylation of KNL-2, the *C. elegans* homologue of M18BP1, is important for mitotic chromosome condensation in nematodes^66^. Mutation of CDK1 phosphorylation sites in KNL-2 resulted in reduced levels of condensin II on mitotic chromosomes and chromosome condensation defects. As the location of KNL-2 phosphorylation sites is not conserved between nematodes and vertebrates, the role of KNL-2 in mitotic chromosome condensation was proposed to be nematode specific. Our finding of a direct association between M18BP1 and condensin II, and of its regulation by CDK1 phosphorylation, however, strongly argues for a conserved evolutionary mechanism that exploits a single protein for multiple functions – centromere maintenance and condensin II regulation. Importantly, these two functions, although confined within a single protein, appear to be at least in part independent, as the M18BP1_4A_ mutation does not affect CENP-A loading.

M18BP1 binding to condensin II directly competes with the negative regulator of condensin II, MCPH1. Competition occurs through similar linear motifs in M18BP1 and MCPH1, which bind to a common pocket in the CAP-G2 subunit. Interestingly, an analogous regulatory mechanism has been proposed for the cohesin complex. Regulatory factors harbouring “YxF” motifs compete for a common binding pocket on the cohesin complex to allow cohesin to either build loops or hold together the sister chromatids^67,68^. This binding pocket on cohesin is located on the SA1 or SA2 subunits, paralogs of CAP-G2. Similarly, yeast condensin and human condensin I may have similar “binding hubs” in the paralogue subunits to CAP-G2^69,70^. Thus, distinct but related SMC complexes may deploy similar mechanisms to regulate their activity spatially and temporally. Collectively, these observations suggest a universal principle of regulation for crucial genomic events across the eukaryotic tree of life.

## MATERIAL AND METHODS

### Cell culture

All cells were cultured at 37 °C in a 5% CO_2_ atmosphere. HAP1 cells were cultured in Iscove’s modified Dulbecco’s medium (Invitrogen) with 10% FBS (Clontech) and 1% penicillin-streptomycin (Invitrogen). HeLa Flip-In T-REx cells were maintained in DMEM medium (Pan Biotech) supplemented with 10% tetracycline-free FBS (Pan Biotech), 50 μg/mL Penicillin/Streptomycin (PAN Biotech), and 2 mM L-glutamine (PAN Biotech). RPE1 cells expressing endogenously tagged M18BP1-mNeonGreen were maintained in DMEM/F-12 medium (Pan Biotech) supplemented with 10% tetracycline-free FBS (Pan Biotech), and 50 μg/mL Penicillin/Streptomycin (PAN Biotech).

### Genome editing

Endogenous mutations were generated in HAP1 cells using CRISPR-Cas9 technology. Oligonucleotides encoding guide RNAs targeting M18BP1 (5’- TTGTACTGAAAAAATCATCA-3’) were cloned into pX459-v2 and co-transfected using FuGENE 6 (Promega) with pUC19 containing a 1528 base pair stretch containing the mutated sequence of the locus of interest and homology arms. To select clones, cells were treated with 2 μg/μl puromycin for 2 days before picking colonies. Mutations were validated by Sanger sequencing the isolated genomic DNA. Endogenously tagged CAP-H2-Halo-3xHA cells were generated using guide RNAs targeting the C-terminus of CAP-H2 (5’- CGGTGCTCCCCACTCAGGGC-3’) in pX330 and a repair template in pUC19. These cells were used as a parental cell line to generate M18BP1 mutant lines described above.

CAP-H2 knockout HAP1 cells were generated as previously described^42^. CAP-H knockout HAP1 cells were generated by insertion of a blast cassette using CRISPR-Cas9 technology. Oligonucleotides encoding guide RNAs targeting CAP-H (5’- GGACTCTGTATACATCGGCA-3’) were cloned into pX459-v2. Knockout cell lines were confirmed by PCR genotyping, Sanger sequencing and immunoblotting.

HeLa CENP-A-SNAP cell lines co-expressing mNeonGreen-CAPH2 and mCherry- M18BP1 variants were generated by transfecting HeLa CENP-A-SNAP Flp-In T-REx cells^29^ with pcDNA5 plasmids and pOG44 plasmid according to the protocol previously described^71,72^.

RPE1 M18BP1-mNG-FKBP-V knock-in cell line was generated via electroporation of gRNA-Cas9 ribonucleoproteins (RNPs) as previously described^73^. Briefly, 2x10^5^ parental hTERT-RPE1 Flp-In TRex (a gift from Johnathon Pines) were electroporated with 200 ng of donor DNA, 120 pmol Cas9, 1.5 μl Alt-R^®^ CRISPR-Cas9 crRNA (AATGAGAAAATATGATTCCT,100 μM, IDT), 1.5 μl Alt-R^®^ CRISPR-Cas9 tracrRNA (100 μM, IDT) and 1.2 μl of Alt-R^®^ Cas9 Electroporation enhancer (100 μM, IDT) using P3 Primary Cell Nucleofector^®^ 4D Kit and Nucleofector 4D system (Lonza). After electroporation, cells were treated with 1 µM NU7441 for 48 h. Individual clones were isolated using FACS sorting. Genomic DNA of monoclonal cell lines was extracted and the correct in-frame knock-in was confirmed by Sanger sequencing of PCR products spanning the cut and insertion sites (Forward primer: TGCTCTCAAGTGGACAGACT; Reverse primer: ACCTCTGTCATCCTTCTCACCT).

### Chromosome spreads

Cells were treated with 250 ng/μl nocodazole for 1.5 hours and mitotic cells were collected by shake-off. Cells were incubated with 75 mM KCl for 10 minutes at 37°C. Cells were pelleted and resuspended in fixative (methanol: acetic acid, 3:1) and incubated for 10 minutes at room temperature. Finally, cells were resuspended in fixative with DAPI and dropped onto microscope slides before mounting with Prolong Gold (Invitrogen). Spreads were imaged using the Metafer system (Metasystems). Images were then randomized using a homemade ImageJ macro and blindly assigned a phenotype.

### Immunofluorescence

Hap1 cells were grown on coverslips and then incubated with 400 nM Janelia Fluor HaloTag ligand 646 (Promega) for 30 minutes then washed three times with media and incubated in fresh media for 30 minutes. Cells were fixed with 4% formaldehyde for 10 minutes then permeabilised with PBS containing 0.15% Triton x100 before blocking in PBST (PBS with 0.05% Tween 20) containing 1% BSA for 1 hour. The following antibodies were used: CENP-A (MA1-20832, Invitrogen, 1:1000). Secondary antibodies (Molecular Probes, Invitrogen) were used at 1:1000 for 1 hour. Coverslips were mounted with Prolong Glass Antifade Mountant with NucBlue stain (Invitrogen).

For depletion experiments and compensation assays, HeLa cells were seeded on coverslips and transfected with Lipofectamine RNAiMAX (Invitrogen), the appropriate siRNA oligo (20 nM M18BP1 - 5’-GAAGUCUGGUGUUAGGAAAdTdT-3’; 50 nM CAPG2-5’- CUCUGAAGUUCGAUCAAAUdTdT-3’; 200 nM MCPH1-5’-CUCUCUGUGUGAAGCACCUdTdT-3’) in serum-Free Opti-MEM medium (Gibco). The transfection controls were set up as above but without adding the siRNA oligos. All conditions were fixed 48 hours from transfection. Exogenous protein expression was induced 24h after transfection by adding Doxycycline (Sigma) to the media at a concentration of 50 ng/ml and induction was performed for 24h. To enrich metaphase cells, 9 µM RO-3306 (Merck Millipore) was added to the media 24h after transfection and cells were incubated for 22 h. Then, drugs were washed out with regular DMEM media and released in media containing 50 ng/ml Doxycycline (Sigma-Aldrich) and 10 µM MG132 (MilliporeSigma). Cells were incubated in the new media for 2h before fixation to allow the enrichment of metaphase states.

For the study of the localisation of M18BP1 during the cell cycle, HeLa or RPE1 M18BP1- mNG-FKBP-V knock-in cells were asynchronously grown on coverslips and fixed 48h after seeding. The different cell cycle states were identified by using the appropriate markers in immunostaining.

To assess the deposition of the new pool of CENP-A in early G1, HeLa cells were seeded into 12-well dishes and treated with siRNA as described above. 24h hours after transfection, cells were exposed to media containing 50 ng/ml Doxycycline (Sigma-Aldrich) and 2 mM Thymidine (Sigma-Aldrich) and were incubated for 16h to enrich for cells in G1/S states. The following morning, the drugs were washed out, existing CENP-A-SNAP proteins were labelled for 30 min using 10 µM SNAP-Cell Block (NEB) and cells were exposed to media containing 5 µM S-trityl-L-cysteine (STLC - Sigma-Aldrich) and were let progress through S, G2 and arrest in mitosis for 7h. Then, the newly produced CENP-A-SNAP pool was labelled by exposing the mitotic cells to media containing 3 µM SNAP-Cell 647 (NEB) and 5 µM STLC (Sigma-Aldrich) for 30 min. Once the labelling was completed, the mitotic cells were collected by shake-off, the drugs were washed out and cells were plated in 24-well dishes containing coverslips. Cells were allowed to attach to the bottom of the wells and exit mitosis for 2.5h, then were fixed and immunostained.

HeLa and RPE1 cells were fixed using ice-cold MeOH for 1 min, then washed and rehydrated 3 times for 5 min with PBS + 0.1% Tween 20 (PBST). Cells were blocked for 20 min with PBST + 5% BSA (Pan Biotech) and then were incubated in wet chambers overnight in primary antibodies. The following morning, coverslips were washed 3 times for 5 min with PBST and then incubated at room temperature in secondary antibodies for 30 min. Finally, the coverslips were washed 3 times for 5 min with PBST and mounted on microscope slides using Mowiol (EMD Millipore) as the mounting agent. The primary antibodies used for these experiments are the following: CAPD3 (A300-601A, Bethyl Laboratories, 1:500), CENP-C (Musacchio Lab, 1:1000), CREST (SKU:15-234, Antibodies Incorporated, 1:2000), CyclinB1 (sc-245, Santa Cruz Biotechnology, 1:1000), CyclinB1 (ab32053, Abcam, 1:1000), M18BP1 (Musacchio lab, 1:1000), PCNA (2586S, Cell Signalling Technology), α-Tubulin (T9026, Sigma Aldrich, 1:500). DNA was visualised using DAPI stain (Sigma, 1:10,000). Fluorescently conjugated secondary antibodies were purchased from Jackson ImmunoResearch Laboratories and used in 1:1,000 dilution. All antibodies were diluted in PBST + 1% BSA.

### Chromatin purification

For the immuno-precipitation coupled to mass-spectrometry experiments of Figure 1D and Figure S1, chromatin was purified either from HeLa FlpIn T-REx cells expressing endogenous CENP-A-SNAP and co-expressing Doxycycline inducible GST-EGFP-M18BP1 and mCherry- MIS18α or from HeLa T-Rex FlpIn cells co-expressing Doxycycline inducible EGFP and mCherry as control. Cells were seeded in T175 flasks and arrested for 18 hours in media containing 10 µM STLC (Sigma-Aldrich) and 50 ng/ml Doxycycline (Sigma-Aldrich) to arrest them in mitosis and induce the expression of the exogenous proteins. The arrested cells were subsequently treated with 500 nM Reversine (Cayman Chemical Company), 9 µM RO-3306 (Merck Millipore) and 10 µM Roscovitine (AdipoGen Life Sciences) to induce mitotic exit. After 3 h cells were harvested by trypsinisation, pelleted and flash-frozen in liquid Nitrogen.

For the immuno-precipitation experiment of Figure 1E, chromatin was purified either from HeLa FlpIn T-REx cells expressing endogenous CENP-A-SNAP and co-expressing Doxycycline inducible EGFP-M18BP1 and mCherry-MIS18α or from HeLa T-Rex FlpIn cells co-expressing Doxycycline inducible EGFP and mCherry as control. Cells were seeded in 10 cm dishes and arrested for 24h in media containing 5 µM STLC (Sigma-Aldrich) and 50 ng/ml Doxycycline (Sigma-Aldrich) to arrest them in mitosis and induce the expression of the exogenous proteins. Cells were harvested by mitotic shake-off, pelleted and flash-frozen in liquid Nitrogen.

Chromatin was purified following a modified version of the protocol from^74^. The pellets were resuspended in 5 volumes of buffer containing 20 mM HEPES pH 7.7, 200 mM KCl, 5 mM MgCl_2_, 1 mM TCEP and supplemented with protease inhibitor cocktail (Serva). Cells were lysed by hypotonic swelling for 2 minutes at room temperature and for 10 minutes on ice. At the end of the incubation, 0.1% NP-40 was added to the tube and the content was mixed by inverting. The lysates were spun at 500g for 10 minutes at 4°C. The cytoplasmic fraction contained in the supernatant was removed and flash-frozen in liquid Nitrogen. The pellet, containing low-purity nuclei, was washed twice with washing buffer (buffer as above supplemented with 0.1% NP-40) and spun at 500 g for 10 minutes at 4°C to remove cytoplasmic impurities. The pellet was resuspended in 5 volumes of washing buffer and sonicated in a Bioruptor Plus (Diagenode) for 5 cycles of 30 seconds ON, and 30 seconds OFF at 4°C. 0.1 µL of Benzonase nuclease (Merck) was added per 200 µL of sample and samples were incubated at 37°C for 5 minutes, followed by incubation at 4°C for 1 hour on a rotor. At the end of the incubation, NaCl was added to a final concentration of 420 mM and salt extraction of the chromatin was performed for 1 hour at 4°C on a rotor. Finally, the samples were centrifuged at 18,000 g for 20 minutes at 4°C. The supernatant containing the solubilised, sheared chromatin fraction was flash-frozen in liquid Nitrogen and stored at -80°C.

### Co-immunoprecipitation

For the experiment of Figure 2H, cells were pelleted, washed with cold PBS and then lysed for 4 hours on rotation at 4°C in TNEN buffer (50 mM Tris pH 7.5, 1 mM EDTA, 150 mM NaCl, 0.1% NP-40, Protease inhibitors (Roche) and phosphatase inhibitors (Sigma, 1:100) supplemented with Ambion DNaseI (Invitrogen, AM2222, 1:100) and Benzonase nuclease (Millipore, 70746, 600 U/ml). After centrifugation at 12,000 rpm for 10 minutes at 4°C, the supernatant was collected and two volumes of TNENG (TNEN buffer supplemented with 10% glycerol) were added. Lysates were quantified by Bradford assay. Protein lysate and Anti-HA magnetic beads (Pierce 88837) were mixed at a ratio of 10:1 (μg:μl) and incubated overnight at 4°C on rotation. Beads were washed three times with wash buffer (50 mM Tris pH 7.5, 1 mM EDTA, 150 mM NaCl, 0.1% NP-40) and proteins were eluted with Laemmli buffer at 95°C for 10 minutes. Co-immunoprecipitation was assessed by immunoblotting.

For the experiment of Figure 1E, 2 mg of purified chromatin from each sample were mixed with 25 µL of GFP-Trap Agarose (ChromoTek) pre-equilibrated with a buffer containing 20 mM HEPES pH 7.7, 20 mM KCl, 300 mM NaCl, 5 mM MgCl_2_ and 0.01% Tween-20. The tubes were incubated for 1 hour at 4°C with gentle rotation. Then, the tubes were centrifuged for 5 minutes at 2500 g. The supernatant containing the unbound fraction was removed, and the beads were washed 3 times with 250 µL of buffer, with a 5-minute centrifugation at 2500 g between each wash. After the final wash, the proteins were eluted in 50 µL of 2X Laemmli buffer. 10% of the volume of the purified chromatin used for each sample was taken as input controls. Samples were boiled at 95°C for 5 minutes and results were assessed by immunoblotting.

### Immunoblotting

Cells were lysed in RIPA buffer (10 mM Tris-Cl (pH 8.0), 1 mM EDTA, 0.2 mM EGTA, 1% Triton X-100, 0.1% sodium deoxycholate, 0.1% SDS and 140 mM NaCl) and quantified by Bradford. 20 μg protein was run on 4-12% Bis-Tris polyacrylamide gels and transferred to PVDF membranes. Membranes were blocked with TBST (TBS and 0.1% Tween 20) and 5% milk for 1 hour. Primary antibodies were diluted in TBST/Milk as follows: CAP-D3 (A300- 604A, Bethyl Laboratories, 1:500), CAP-G2 (A300-605A, Bethyl Laboratories, 1:500), CAP-H (NBP1-32573, Novus Biologicals, 1:1000), CAP-H2 (A302-275A, Bethyl Laboratories, 1:500 and 1:1000), GFP (Musacchio Lab, 1:500), Histone H3 (ab10799, Abcam, 1:500), MCPH1 (4120, Cell Signalling Technologies, 1:2000), M18BP1 (Musacchio lab, 1:300), Tubulin (T5168, Sigma Aldrich, 1:50,000), α-Tubulin (T9026, Sigma Aldrich, 1:10,000).

### Affinity Purification LC-MS/MS and data analysis

Triplicates of immunoprecipitated GFP-M18BP1 and GFP alone were directly digested on beads^75^. The peptides were subsequently separated on an UltiMate^TM^ 3000 HPLC System (ThermoFisher Scientific) using a 90 min gradient from 5-60% acetonitrile with 0.1% formic acid and directly sprayed via a nano-electrospray source in a quadrupole Orbitrap mass spectrometer (Q Exactive^TM^, Thermo Fisher Scientific^TM^)^76^. Data was acquired in a data-dependent mode acquiring one survey scan (MS scan) and subsequently 15 MS/MS scans of the most abundant precursor ions from the survey scan. The mass range was set to m/z 300 to 1600 and the target value to 3x10^6^ precursor ions with a 1.4 Th isolation window. The maximum injection time for purified samples was 28 msec. MS scans were recorded with a resolution of 60.000 and MS/MS scans with 15.000. Unassigned precursor ion charge states and singly charged ions were excluded. To avoid repeated sequencing, already sequenced ions were dynamically excluded for 30 sec. The resulting raw files were processed with the MaxQuant software (version 1.6.14) using N-terminal acetylation, oxidation (M) as variable modifications and carbamidomethylation (C) as fixed modification. Label-free quantification was enabled. A false discovery rate cut-off of 1% was applied at the peptide and protein levels and as well on the site decoy fraction^77^. The MaxQuant proteingroups.txt output table was then further processed in Perseus (version 1.6.50)^78^. Contaminants and reverse hits were removed. To obtain a list of confident interaction partners proteins were filtered for quantification in at least 2 replicates out of 3 replicates. Missing values were imputed with a downshift of 1.8 and a width of 0.3. The cut-off lines for the volcano plot were set to p-value < 0.01 and S0=2.

### Synthetic lethality screen

Synthetic lethality screens were carried out as previously described^79^. In brief, haploid cell lines were infected with gene-trap retroviruses which, when integrated in a disruptive (sense) orientation into gene introns, can create a knock-out. By culturing for 12 days, cells with a gene knockout important for cell viability will be depleted from the population and cells in which the virus integrates in a non-disruptive (anti-sense) orientation will survive. As disruptive and non-disruptive integrations occur at similar frequencies, the ratio of insertions in the surviving population indicates whether a gene is important for cell fitness in the specified genetic background. To determine this ratio, cells were harvested and fixed in fix buffer I (BD biosciences). G1 haploid cells, defined by DAPI intensity, were sorted out by flow cytometry on a BD FACSAria Fusion. Genomic DNA was isolated and LAM-PCR carried out to identify insertion sites by sequencing on a Illumina NovaSeq SP with 100bp single reads. Insertion sites were mapped using standard procedures^79^, with some changes. In summary, the unique reads were aligned against the hg38 human genome using Bowtie, allowing for no more than one mismatch. Subsequently, these reads were assigned to protein-coding genes using the longest open reading frame transcript and excluding overlapping regions that cannot be attributed to a single gene. The count of unique alignments was conducted within intronic regions spanning from the transcription start site to the stop codon. A false discovery rate–corrected binomial test was employed (FDR-corrected P value) to identify the overrepresentation of genes in either the sense or antisense orientation of gene-trap insertions. Additionally, the significance of genes after genetic perturbation (*ΔCAP-H)* compared to wild-type control cells was evaluated using a bidirectional Fisher’s exact test across four independent control datasets.

### Microscopy and image analysis

Cells from Figure S3.1A, S3.3A, S3.4A, S5.3A were imaged on a DeltaVision Elite deconvolution microscope (GE Healthcare, UK), equipped with an IX71 inverted microscope (Olympus, Japan), a UPLSAPO x100/1.40NA oil objective (Olympus) and a pco.edge sCMOS camera (PCO-TECH Inc., USA). Cells from Figure 5A, 5C, S3.2A, S5.1C were imaged on a spinning disk confocal device on the 3i Marianas system equipped with an Axio Observer Z1 microscope (Zeiss), a CSU-X1 confocal scanner unit (Yokogawa Electric Corporation, Tokyo, Japan), x100/1.4NA oil objective (Zeiss), and Orca Flash 4.0 sCMOS Camera (Hamamatsu). All the images were acquired as z-sections at 0.20 μm.

HAP1 cells from Figure 3A were imaged on an AxioObserver Z1 (Zeiss) microscope with an x63 oil immersion objective and z stacks of 0.2 μm. Images of HAP1 cells were converted to maximum intensity projections for analysis. The acquired images were converted into sum intensity projections, exported, and converted into 8-bit using ImageJ^80^. Quantification of the chromatin signal was performed on Fiji using a script for semiautomated processing. Briefly, Regions of interest (ROIs) were established by segmenting the chromatin DAPI signal via Otsu thresholding^81^. Applying those ROIs to the respective fluorescent channels yielded mean protein signal intensities. Correction for background intensity was done by subtracting from the mean protein intensity the mean intensity from the border region around the chromatin. This border region was bounded to the inside by the ROI and to the outside by the ROI, which was slightly enlarged by repeated binary dilation. Absolute signal intensities were calculated by multiplying background-corrected mean intensities with chromatin ROI area. Data was normalised using Excel (Microsoft) and plotted using GraphPad Prism (GraphPad) software. Data was visualised as violin plots in Prism 8. For each sample, the median value of each repeat was superimposed to the violin plots as described in the SuperPlots methodology^82^. The figures were arranged using Adobe Illustrator software.

Inter-kinetochore distances were measured manually in Fiji software. DAPI and CREST channels from deconvolved images were used to identify kinetochores of the same bi-oriented sister chromatids pair. Measurements were performed in Z-stacks where both the sister kinetochores were visible, and the distance between the CREST signals was measured using a straight line. 10 inter-kinetochore distances were randomly measured for each cell and the average value was used in the final plotting.

### Statistical analysis

Statistical analysis was performed in Prism 8. All cell biology experiments were repeated at least 3 times to attain statistical significance. We used the Mann-Whitney test for experiments comparing two conditions, and the Kruskall-Wallis test coupled with Dunn’s multiple comparisons for experiments comparing 3 or more conditions.

### Protein expression and purification

Condensin I and II pentamers were purified as described in^43^ and condensin II tetramer, MBP- MCPH1_1-195_, MBP-MCPH1_196-435_, and MBP-MCPH1_1-435_ were purified as described in^21^.

M18BP1_873-1132_ and M18BP1_873-1132_ DDFF to AAAA mutant version were expressed in Rosetta (DE3). Cells were grown in Terrific Broth media at 37 degrees. Upon reaching OD of about 4, cells were induced for 2 hrs at 30 degrees, then centrifuged and flash frozen in liquid nitrogen. Cells were resuspended in HEPES pH 7.5, NaCl 200 mM, glycerol 10% and DTT 2mM, and lysed by sonication on ice at 60% amplitude, 30 second on and 30 seconds off for a total of 5 minutes. After centrifugation the supernatant was injected on a His Trap column and eluted with increasing concentration of Imidazole. The His tag purification was followed by a MBP affinity purification and finally a size exclusion column. Phosphorylation of M18BP1, was obtained by treating the purified protein over night at 4 degrees with CDK1:Cyclin-B:CKS1 complexes in presence of 2 mM ATP and 10mM MgCl_2_.

### Sulfo-SDA crosslinking reaction and peptide preparation for crosslinking MS

Condensin II - M18BP1 complex at 1.5 uM was incubated with AMPPNP at 1mM concentration and reacted with sulfo-NHS-diazirine (sulfo-SDA, Thermo Scientific) at 0.5/1/1.5mM. The crosslinked protein material was then separated on a 4-12% bis-tris SDS-PAGE gel (life technologies) and the bands corresponding to the crosslinked protein complex were excised and processed by in-gel digestion^83^. Briefly, the proteins were reduced with 20mM dithiothreitol (DTT, thermo scientific) and alkylated with 55mM iodoacetamide (IAA, merck millipore) prior to digestion with trypsin (Pierce). Peptides were then recovered and desalted with C18 StageTips (Empore) and crosslinked peptide pairs were enriched by size exclusion chromatography (SEC)^84^ using a superdex 30 increase column (Cytiva) equilibrated with 30% acetonitrile, 0.1% trifluoroacetic acid. 50ul fractions were collected and early eluting fractions were taken for LC-MS.

### Sulfo-SDA Crosslinking MS acquisition

Approximately 1ug of peptides of each fractionwere injected for each liquid chromatography-mass spectrometry (LC-MS) acquisition. The LC-MS platform consisted of a Vanquish Neo system (ThermoFisher Scientific) connected to an Orbitrap Eclipse Tribrid mass spectrometer (ThermoFisher Scientific) equipped with a FAIMS Pro Duo device operating under Tune 3.5.3886. Mobile phases consisted of 0.1% v/v formic acid in water (mobile phase A) and 0.1% formic acid in 80% acetonitrile/water v/v (mobile phase B). Samples were dissolved into 4% mobile phase B. The FAIMS Pro Duo device was set to standard resolution with a carrier gas flow of 4.6L/min. The samples were separated on an EASY-Spray PepMap Neo column (75 µm x 50 cm) (ThermoFisher Scientific). Peptides were separated on 110 minute methods designed to match the hydrophobicity of the various SEC fractions, with linear separation gradients from 20%-40%B in 77 minutes (earliest fraction, most hydrophobic), down to 11%B to 35%B (latest fraction, least hydrophobic).

MS1 spectra were acquired with a resolution of 120,000 and automated gain control target set to 250% and 50ms maximum injection time. Source RF lens was set to 35%. Dynamic exclusion was set to single count in 60 seconds. The duty cycle was set to 2.5 seconds. A precursor charge filter was set to z=3-7. Precursors were selected based on a data-dependent decision tree strategy prioritizing charge states 4-7 and subjected to stepped HCD fragmentation with normalized collision energies of 20, 27, 30^84^. MS2 scans were acquired with a normalized gain control target of 750% with maximum injection time of 250ms and an orbitrap resolution of 60,000. For each SEC fraction, multiple injections were carried out with multiple FAIMS control voltages, with the first 3 injections being performed at -45V, -55V and -65V separately, and, if possible, a final injection with a 2 CV combination of -40/-75V each with a duty cycle of 15 seconds .

### Sulfo-SDA Crosslinking MS data analysis

Raw files were converted to mgf format using ProteoWizard MSconvert (version 3.0.22314). A recalibration of the MS1 and MS2 m/z was conducted based on high-confidence (<1% false discovery rate) linear peptide identifications using xiSEARCH (version 1.6.745). Crosslinking MS database search was performed in xiSEARCH (version 1.7.6.7)^85^ on a database comprising human condensin II, M18BP1, contaminants from protein purifications, and common mass spec contaminants derived from MaxQuant^77^ searched with 4 missed tryptic cleavages. Precursor mass error tolerance was set to 3ppm and MS2 error tolerance to 5ppm. The search included methionine oxidation, asparagine deamidation, SDA loop link (+82.04186484Da), hydrolized SDA (+100.0524) as variable modifications. Site-specific phosphorylation at T993 of M18BP1 was defined as a variable modification. The SDA crosslinker was defined as cleavable^86^. The search was set to account for noncovalent gas-phase associations. Prior to FDR estimation, search results were filtered to only include peptide spectra matches with at least 2 crosslinker-containing fragments on both peptides. Results were filtered to 5% FDR at the residue pair level and 10% at the protein pair level using xiFDR (version 2.1.5.2) and the “boost” feature to optimize thresholds at the lower error levels was enabled to maximise heteromeric crosslinks. Results were exported in mzIdentML format and uploaded to xiview.org for visualization. Pseudobonds files were downloaded and visualized on the structural model using chimera X version 1.6.1 (Figure S2.5).

### Analytical size exclusion chromatography

Condensin II at final concentration of 10 μM was mixed with M18BP1_873-1132_ (WT or DDFF to AAAA mutant version) at final concentration 20 μM. 50 μL of the reaction mixture were injected onto a 2.4 ml Superose 6 Increase gel filtration column (GE Healthcare) equilibrated in 20 mM TRIS pH 8.5, 150 mM NaCl, 5mM MgCl_2_, 5% (v/v) Glycerol and 2 mM DTT. 50 μL fractions were collected and analysed by SDS–PAGE using 4–12% NuPAGE Bis-Tris gels (Invitrogen). The gels were run in MOPS buffer at 200 V for 45 min and stained with Instant- Blue Coomassie protein stain (Abcam).

### ATPase assay

ATPase activities were measured using the ATPase/GTPase Activity Assay Kit (MAK-113, Sigma-Aldrich) according to the manufacturer’s instruction. The purified protein was diluted to 200 nM with Assay Buffer (20 mM Tris pH 8.5, 50 mM NaCl, 5 mM MgCl_2_, 5% (w/v) glycerol, 2 mM ATP, DTT 2mM). Equimolar concentration of condensin II and M18BP1 were used for the reactions. For the experiments containing DNA, a 10 fold excess of dsDNA to protein was used. 20 μL of the reaction mixture containing the diluted protein was incubated for 30 min at 20 °C. Then, 100 μL of malachite green reagent was added into each reaction well and incubated for 10 min. After that, the absorbance at 620 nm was measured, proportional to the enzyme activity present.

### Fluorescence polarisation protein binding assays

Peptides used in fluorescence polarisation assays were synthesised by Genscript and are shown in Table 2. The concentration of 5-FAM wild-type MCPH1407-422 was determined using the 5-FAM extinction coefficient of 83,000 (cmM)–1 at 493 nm. Non-labelled peptides had TFA removed to less than 1 % and were accurately quantified using Genscript’s amino acid analysis service. All peptides were solubilised in DMSO and diluted to a working concentration in FP assay buffer (20 mM Tris pH 8.5, 200 mM NaCl, 1 mM DTT, Tween20 0.08%, BSA 0.5mg/mL).

Non-phosphorylated MCPH1 FP competition assays were performed using 0.3uM of 5- FAM-labelled MCPH1_407- 424_ peptide, with or without 0.8uM of condensin II pentamer and with 8uM of MBP-MCPH1_1-195_, MBP-MCPH1_196-435_, MBP-MCPH1_1-435_ or MBP-M18BP1_873-1132_ in a total volume of 40 µl in half- area black plates (Constar). The plate was incubated at room temperature for 20 min, before being read with an Omega plate reader (BMG Labtech) at 5 min intervals and monitored to ensure binding had reached equilibrium. Each plate was read three times, and three replicates were performed. The buffer only background was subtracted and mean and standard deviation was calculated.

FP Binding assays with phosphorylated or unphosphorylated 5-FAM-labelled MCPH1_407- 424_ peptides to condensin II was performed using 0.3uM peptides, with increasing concentration of condensin II ranging from 20 nM to 5uM.

FP Competition assays with phosphorylated and unphosphorylated M18BP1 to MCPH1_407-424_ – condensin II complex were performed using 0.3uM of 5-FAM-labelled MCPH1_407- 424_ peptide and 0.6 uM of condensin II pentamer. The complex of condensin II - 5-FAM-labelled MCPH1_407-424_ was incubated with increasing concentration of phosphorylated or unphosphorylated M18BP1 ranging from 30 nM to 8 uM.

The FP binding and competition assays were measured at 25°C in Corning 384-well Black Round Bottom well plates. Three independent measurements were collected and averaged for each point of the binding isotherms. The dissociation constant (K*_D_* and K*_D2_*) values and standard error of the mean were calculated in Graphpad Prism, using all data points from three independent experiments. FP data was fit using equations for direct binding and directly competitive binding^87^.

### Cryo-EM sample preparation and data acquisition

For the preparation of cryo-EM samples (Figure S2.2), purified condensin II and M18BP1 were incubated with 2 mM AMP-PNP for 30 min. The sample was then run on a glycerol (10-25%) and glutaraldehyde (0-0.2%) gradient by ultracentrifugation with a Sw60 rotor set on a G-force of 29k for 16hrs at 4 degrees. After the GRAFIX fixation, the complex was again separated from the aggregates and washed off the glycerol on a Superose 6 Increase 10/300 GL column equilibrated in TRIS pH 8.5, NaCl 150 mM, DTT 2mM, 0.03% beta-glucopyranoside and 0.03% Tween20. The peak fractions corresponding to the right molecular weight were pooled and concentrated. The freshly purified condensin II – M18BP1 complex (OD280 = 0.5) was applied to glow-discharged Quantifoil R1.2/1.3 300-mesh copper holey carbon grids. Grids were blotted for 3.5 s under 100% humidity at 4°C before being plunged into liquid ethane using a Vitrobot Mark IV (FEI). Micrographs were acquired on a Titan Krios microscope (FEI) operated at 300 kV with a Falcon 4i direct electron detector using a slit width of 10 eV. EPU software was used for automated data collection following standard procedures. A calibrated magnification of 105x was used for imaging, yielding a pixel size of 1.2 Å on images. The defocus range was set from -0.5 μm to -2 μm. Each micrograph was dose-fractionated to 50 frames, with a total dose of about 50 e-/Å^2^.

### Cryo-EM image processing

The detailed image processing and statistics are summarized in Figures S2.2, S2.3 and Table 1. Motion correction was performed using the Relion’s own implementation of the MotionCorr2 program^88^, and the CTF parameters of the micrographs were estimated using the CTFFIND program^89^. Most steps of image processing were performed using RELION-3 or RELION-4^90,91^. Initially, particle picking was performed by using the Laplacian-of-Gaussian blob detection method in RELION-3. Good class averages representing projections in different orientations selected from the initial 2D classification were used 3D initial model and 3D classification in RELION. Particles aligning in the best 3D classes were used for Topaz^92^ training and autopicking. Extracted particles were binned 3 times and subjected to 2D classification and 3D classification. The selected classes from 3D classification were subjected to 3D auto refinement followed by Bayesian polishing. Polished particles were used for three rounds of 3D classification with alignment. Finally, we performed 3D classification without local search that yielded to a final set of 49 k particles. Selection of particles was used for 3D auto-refine job and final map was post-processed to correct for modulation transfer function of the detector and sharpened by applying a negative B factor, manually set to -150. A soft mask was applied during post processing to generate FSC curves to yield a map of average resolution of 7 Å. The RELION local resolution procedure was used to estimate local resolution for the map that resulted in a resolution range between 5 and 11 Å. Attempts at Multibody refinement or CryoDRGN^93^ were made to increase local resolution of subunits. However, both programs showed a continuous motion among subunits that could not be resolved in local minima, hence limiting the overall resolution of the map.

**Table 1.** Cryo-EM data collection, refinement, and validation statistics.

| **Table 1. Cryo-EM data collection, refinement, and validation statistics** |  |
| --- | --- |
| **Data collection and processing** | condensin II |
| Magnification | 105,000 |
| Voltage (kV) | 300 |
| Electron exposure e<sup>-</sup>/Å<sup>2</sup> | 50 |
| Defocus range (μm) | 0.8 to 2.0 |
| Pixel size (Å) | 1.2 |
| Symmetry imposed | C1 |
| Initial particle images (no) | 3.7 M |
| Final particle images (no) | 49k |
| Map resolution (Å) | 7 |
| FSC threshold | 1.43 |
| Map resolution range (Å) | 5 to ~ 11 |
| **Refinement** |  |
| Initial model used | Alphafold monomer predictions |
| Model resolution (Å) | 6.6 |
| FSC threshold | 0.143 |
| Map sharpening B factor (Å<sup>2</sup>) | -300 |
| **Model comparison** |  |
| Nonhydrogen atoms | 23364 |
| Protein residues | 2891 |
| **B factors (Å<sup>2</sup>)** |  |
| Protein |  |
| **r.m.s deviations** |  |
| Bond lengths (Å<sup>2</sup>) | 0.005 |
| Bond angles (°) | 1.043 |
| **Validation** |  |
| MolProbity score | 1.64 |
| Clashscore | 8.27 |
| Poor rotamers (%) | 1.77 |
| **Ramachandran plot** |  |
| Favored (%) | 96.85 |
| Allowed (%) | 3.12 |
| Outliers (%) | 0.04 |

**Table 2.** peptides for Fluorescence polarization assays.

| **Table 2 – peptides for Fluorescence polarization assays** |  |
| --- | --- |
| 5FAM-MCPH1 | 5FAM- CGESSYDDYFSPDNLKER |
| MCPH1pS417 | CGESSYDDYF(pSER)PDNLKER |

### Structure and sequence analysis

Sequence conservation predictions were made with ConSurf^94^. AlphaFolds structure prediction was performed using AlphaFold2-multimer was executed using a locally installed version of AlphaFold2^38^. This local version of AlphaFold2-multimer operates with V3 model parameters and utilizes default arguments. The databases employed for the prediction in the step of Multiple Sequence Alignment (MSA) generation are as follows: Uniref30 (UniRef30_2021_03), Uniref90 (downloaded on 04/02/2023), UniProt (downloaded on 03/02/2023), PDB_mmcif (downloaded on 04/02/2023), PDB_seqres (downloaded on 04/02/2023), PDB70 (version 06/09/2014), mgnify (version 05/2022), and BFD.

The multimer prediction was utilized for full length proteins (CAP-G2 with MCPH1 or M18BP1). The highest-ranked structures are displayed in the figures, accompanied by the Predicted Alignment Error (PAE) plot of the highest-ranked model (S2.1 E-F and S4.1). Images for figures were generated using UCSF ChimeraX ^95^.

### Data and code availability

The cryo-EM map of the condensin II-M18BP1 complex has been deposited in the Electron Microscopy Data Bank (EMDB) with accession code EMD-50201. An atomic model of the condensin II-M18BP1 complex has been deposited in the Protein Data Bank (PDB) with accession code 9F5W.

The mass spectrometry proteomics data have been deposited to the ProteomeXchange Consortium (http://proteomecentral.proteomexchange.org) via the PRIDE^96^ partner repository with the dataset identifier PXD051556.

The mass spectrometry crosslinking MS data for condensin II-M18BP1 have been deposited to the ProteomeXchange Consortium (http://proteomecentral.proteomexchange.org) via the PRIDE^96^ partner repository with the dataset identifier PXD051886 and 10.6019/PXD051886.

## ACKNOWLEDGMENTS

We thank Martin Houlard for helpful discussions regarding MCPH1 and Norman Davey for helpful discussion on short linear motifs. We thank P. Swuec and S.Sorrentino of the Cryo-Electron Microscopy Unit of the National Facility for Structural Biology at Human Technopole, for technical support and assistance.

## SUPPLEMENTARY

**Figure S1.1:**
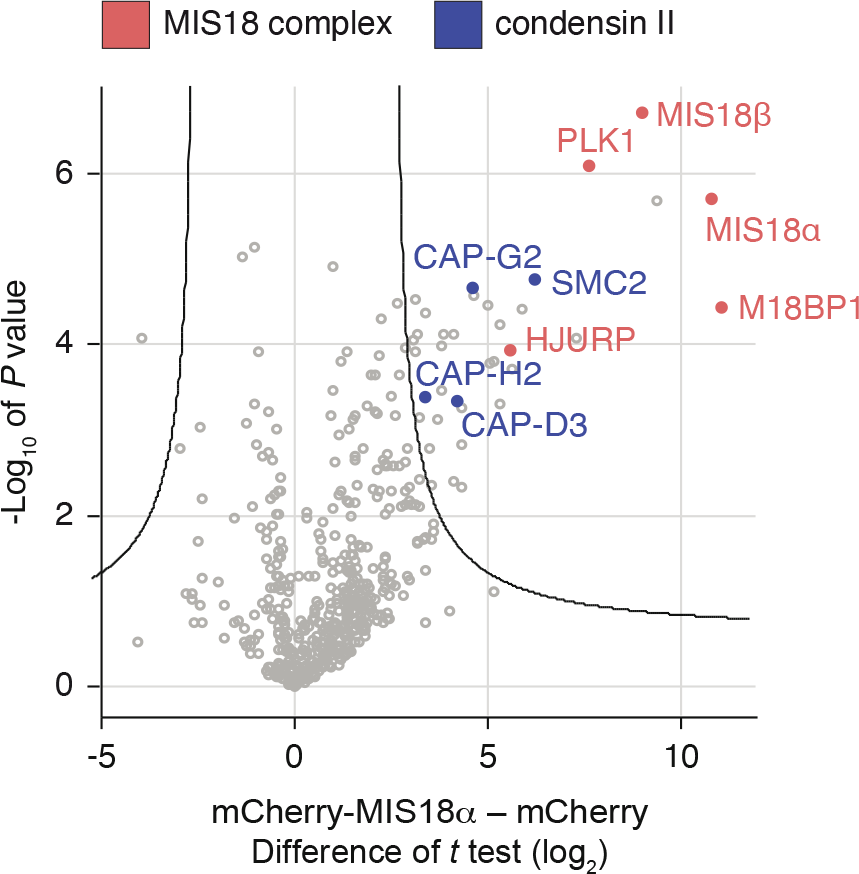
Co-IP of MIS18α from early G1 chromatin identifies condensin II as a possible interactor. Volcano plot shows the chromatin interactome of MIS18α in early G1 phase. Condensin II subunits are marked in light blue whereas the CENP-A deposition machinery components are marked in red.

**Figure S2.1:**
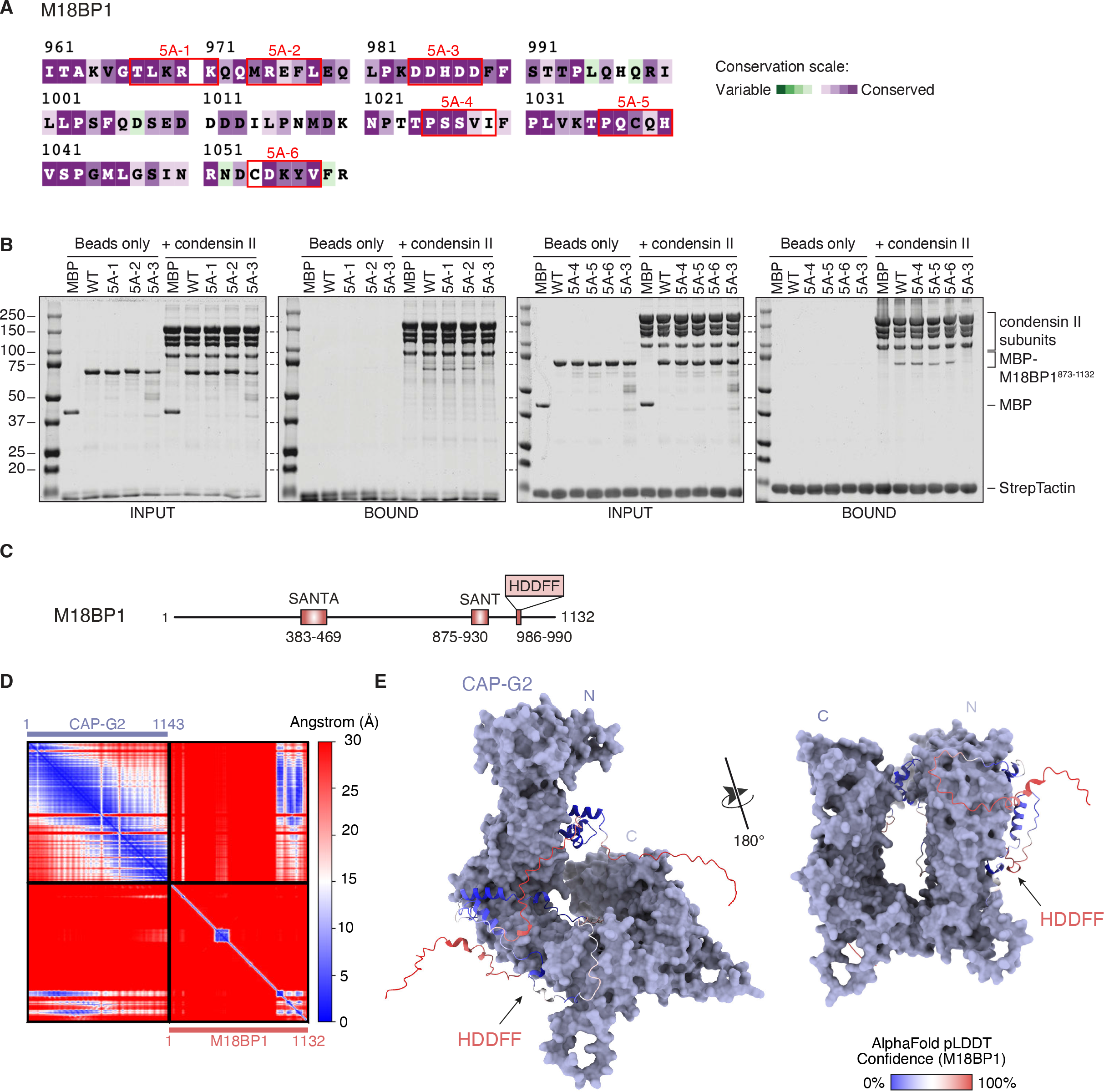
Dissecting the condensin II – M18BP1 interactions. (A) Conservation of the M18BP1 sequence within the region that binds to condensin II. Red rectangles indicate residues selected for mutagenesis to 5x alanine. (B) Streptavidin bead pull down assay with strep-tag condensin II and M18BP1 alanine mutants. (C) Domain schematic of M18BP1. (D) Predicted aligned error (PAE) plot for the interaction between CAP-G2 and M18BP1 blue indicates high confidence and red low confidence in relative positioning (E) Surface representation of AF2 prediction of CAP-G2 (purple) with a ribbon model of M18BP1 (colored by pLDDT score) for the Alphafold Multimer prediction.

**Figure S2.2:**
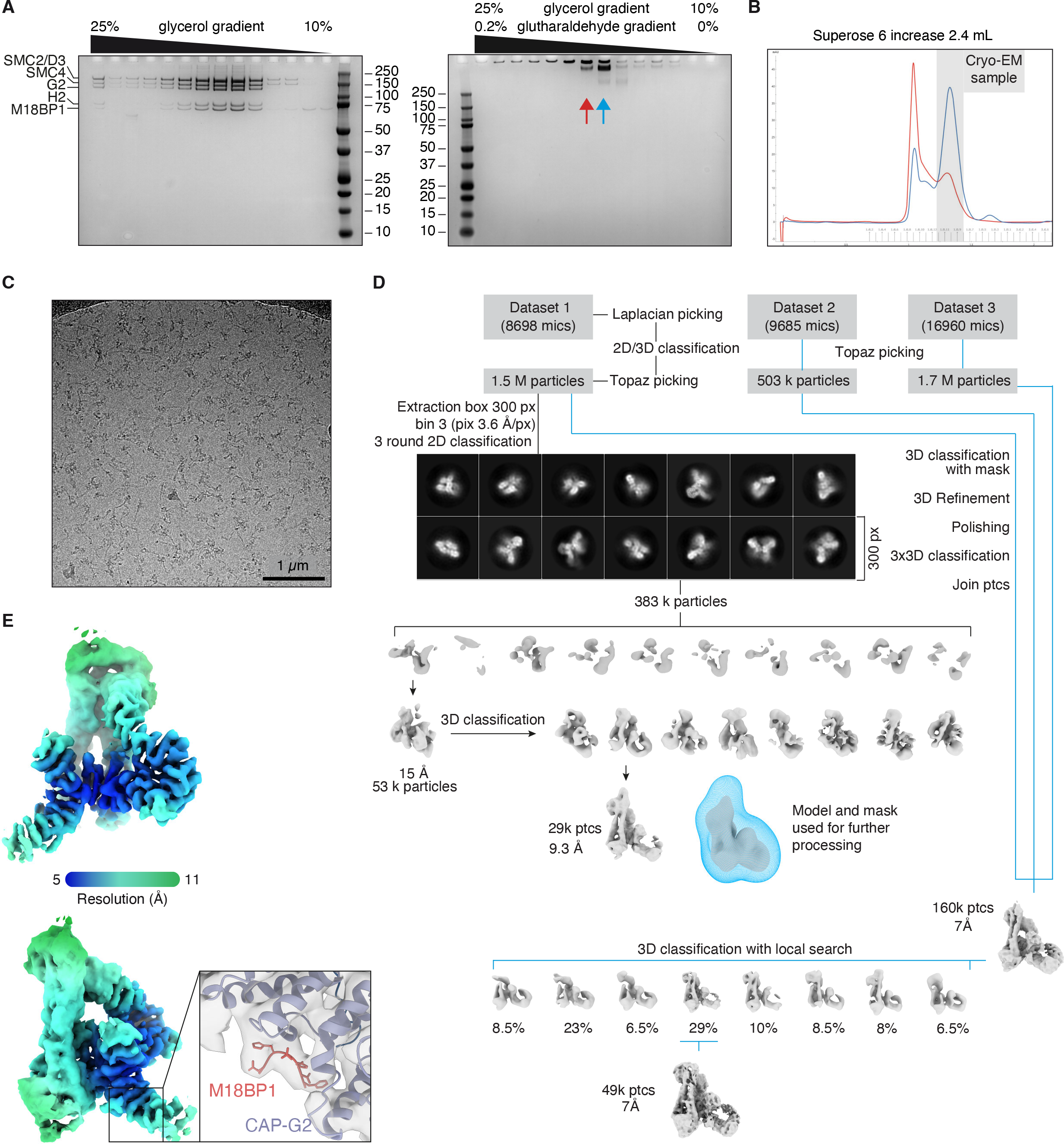
Cryo-EM sample preparation and data processing overview. (A) SDS page of fractions from glycerol gradient in presence (right) and absence (left) of 0.2 % maximum percentage of glutaraldehyde. (B) Size exclusion profiles of gradient fractions 7 (red) and 8 (blue). (C) Representative micrograph from cryo-EM dataset. (D) Cryo-EM processing pipeline with final map colored by resolution (E) and zoom on the extra density for the CAP- G2 (purple) - M18BP1 (red) interaction interface.

**Figure S2.3:**
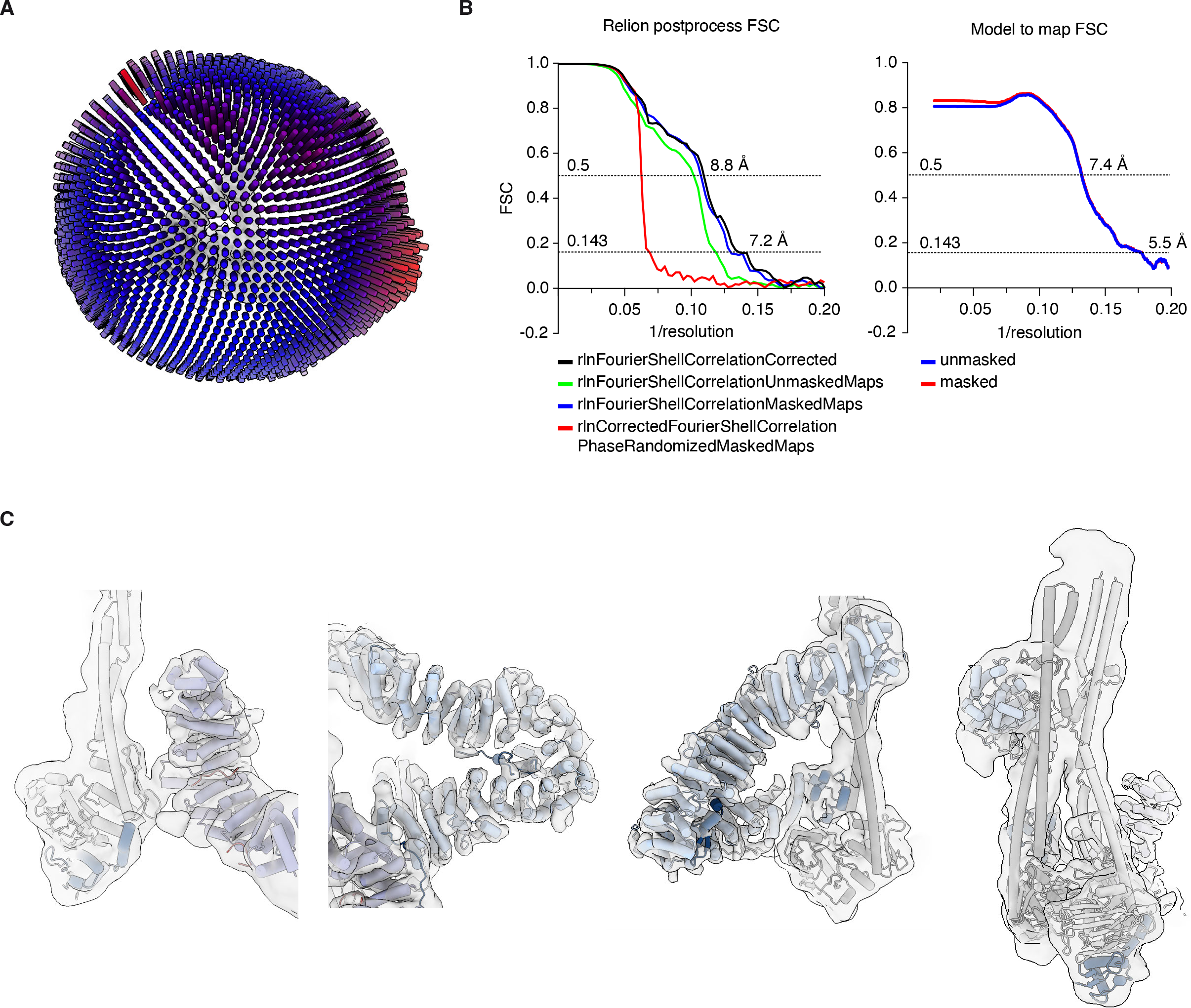
Cryo-EM model and map statistics. (A) Orientation distribution of particles in final cryo-EM map. (B) Fourier Shell Correlation (FSC) curves for raw map obtained with RELION (right) and model to map (left). (C) Details of structural model fit to EM density.

**Figure S2.4:**
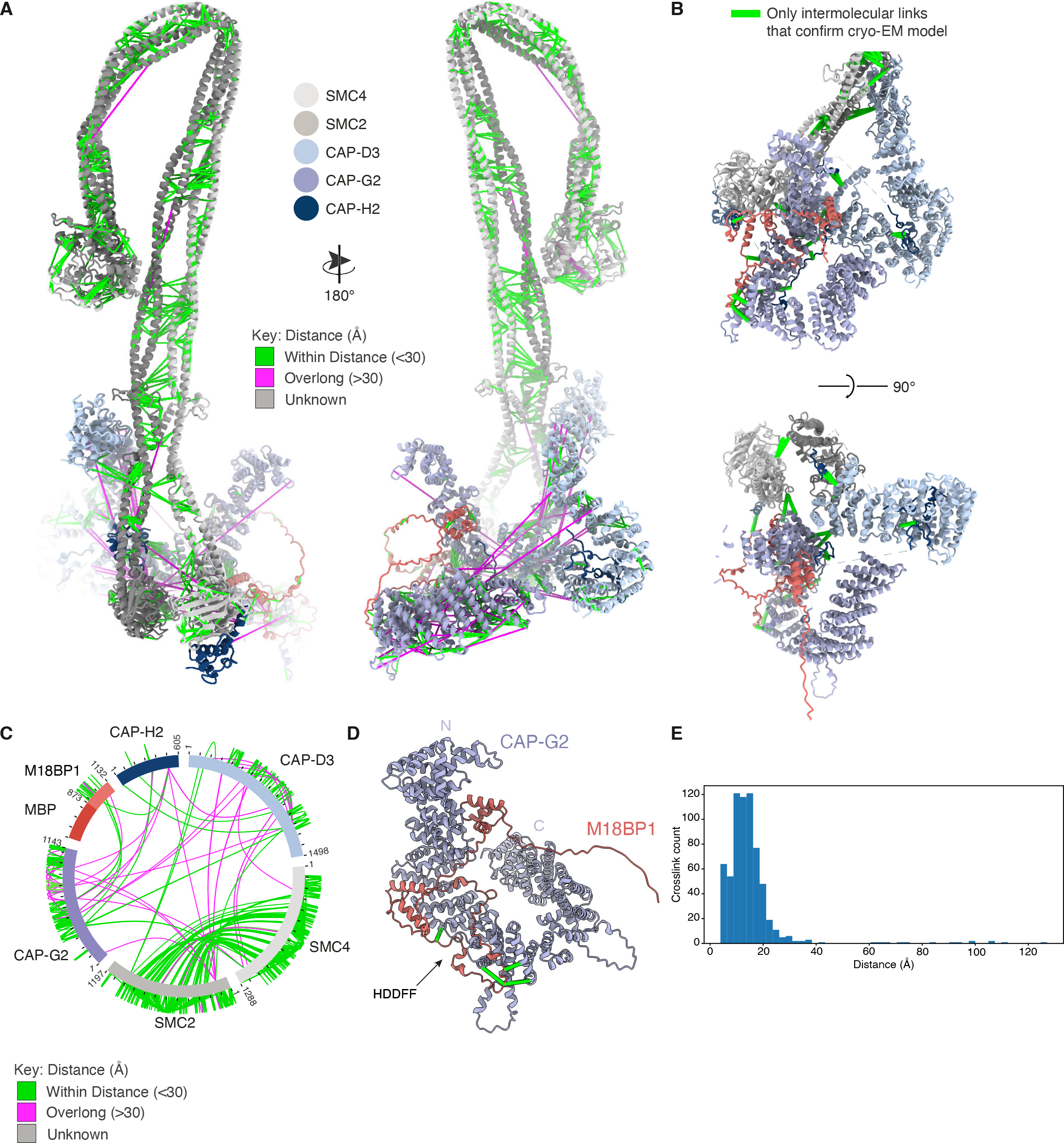
Crosslinking MS analysis. (A) Whole condensin II - M18BP1 complex sulfo-SDA crosslinks. Crosslinks satisfied in the model are coloured in green and overlong crosslinks coloured in magenta. (B) Zoom on core domain of complex with distance respected intermolecular crosslinks coloured in orange. (C) Full crosslinking MS dataset for condensin II - M18BP1 depicted on a circle plot. Crosslinks satisfied in the model are coloured in green and overlong crosslinks are coloured in magenta. Crosslinks in regions missing from the model are coloured in grey. (D) AF2 prediction of NCAPG2 (purple) -M18BP1 (red) with intermolecular crosslinks shown on the structure. (E) Distance distribution plot of whole crosslinking MS dataset.

**Figure S2.5:**
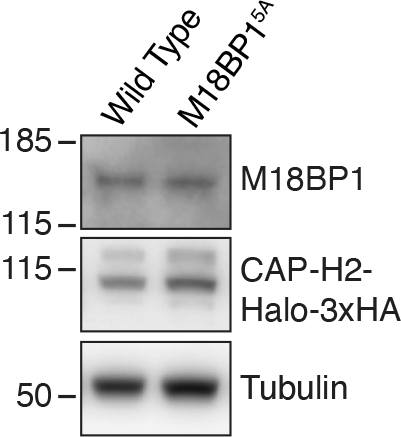
**Control for cell lines used in co-IP (**Figure 2) Levels of endogenously tagged CAP- H2-Halo-3xHA in the indicated cell lines.

**Figure S3.1:**
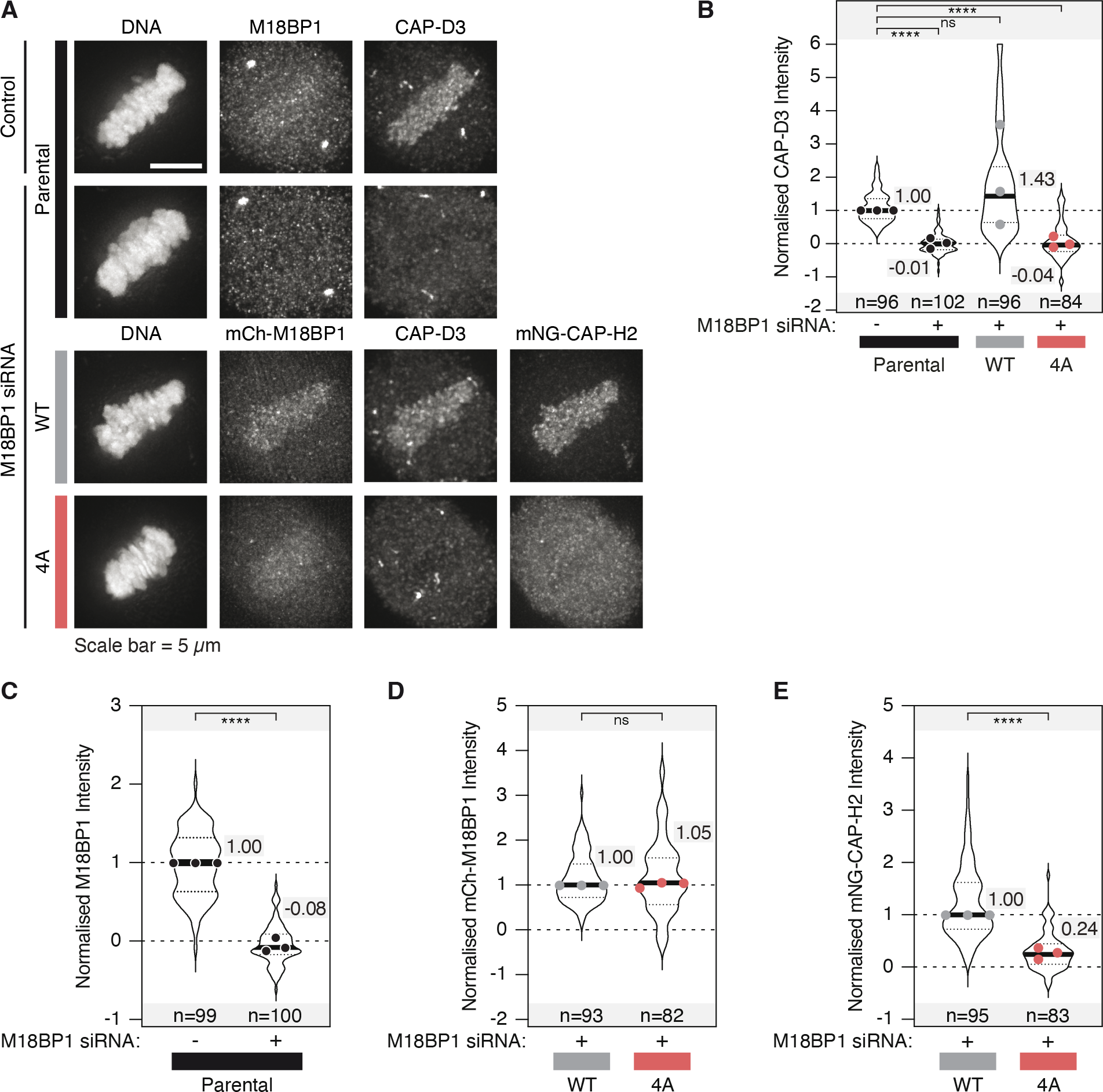
M18BP1 is a major recruiter of condensin II in mitotic HeLa cells. (A) Localisation of endogenous CAP-D3 in HeLa cells conditionally expressing siRNA resistant mCherry- M18BP1 WT or 4A mutant. (B) Quantification of CAP-D3 in the indicated cell lines in A. (C) Quantification of endogenous M18BP1 levels in the depletion controls from A. (D) Quantification of mCherry-M18BP1 levels from A. (E) Quantification of mNeonGreen-CAP- H2 signal from the depletion controls of A. “n” represents the number of cells analysed across three independent experimental repeats. The median of the combined data is shown for each condition and dots show the median of each experimental repeat.

**Figure S3.2:**
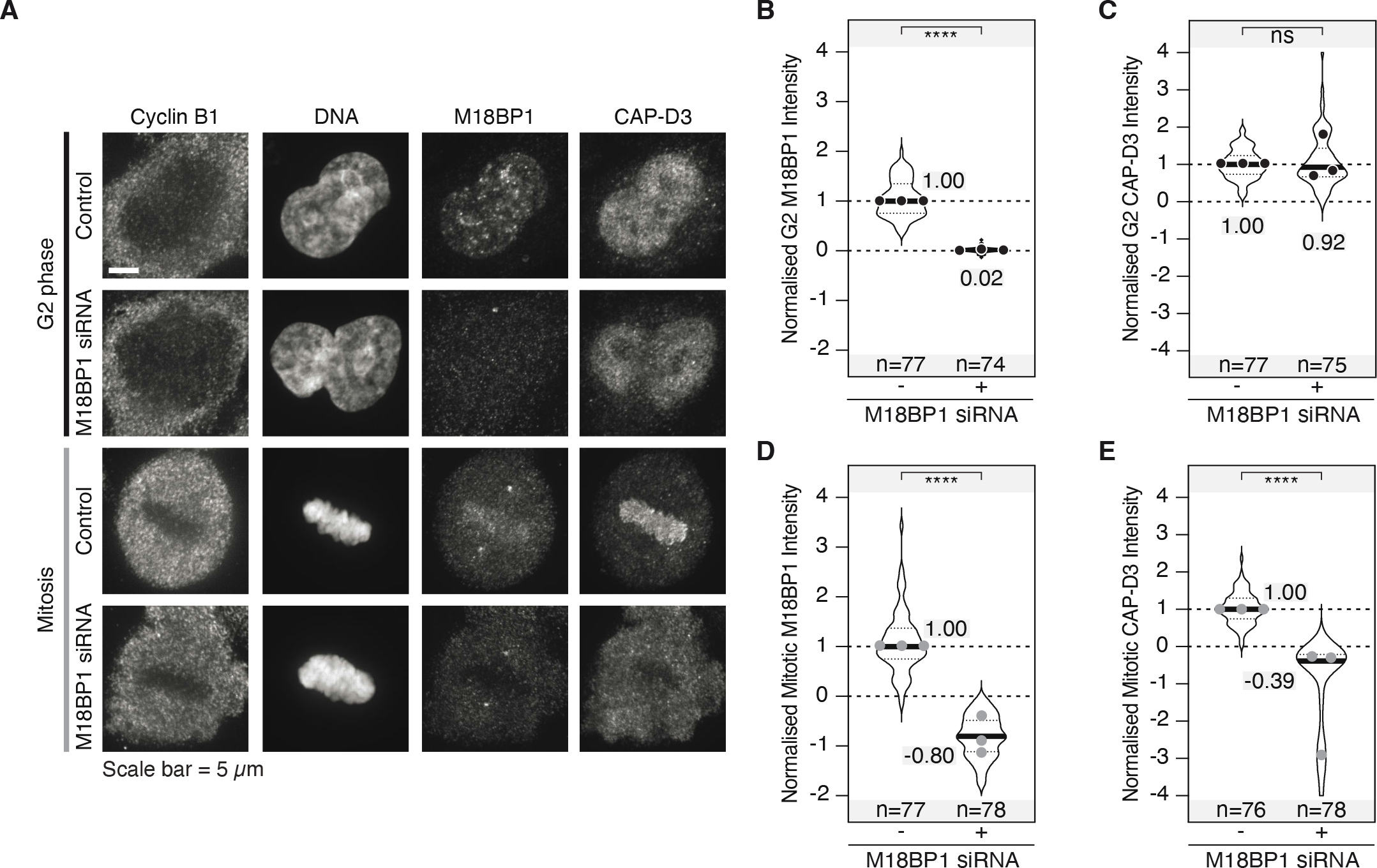
M18BP1 depletion does not affect interphase condensin II localisation. (A) Localisation of CAPD3 in mitotic and G2 cells treated with M18BP1 siRNA. G2 cells were selected based on CyclinB1 levels. (B) Quantification of M18BP1 and (C) CAP-D3 levels in G2 cells from A. (D) Quantification of M18BP1 and (E) CAP-D3 levels in mitotic cells from A. “n” represents the number of cells analysed across three independent experimental repeats. The median of the combined data is shown for each condition and dots show the median of each experimental repeat.

**Figure S3.3:**
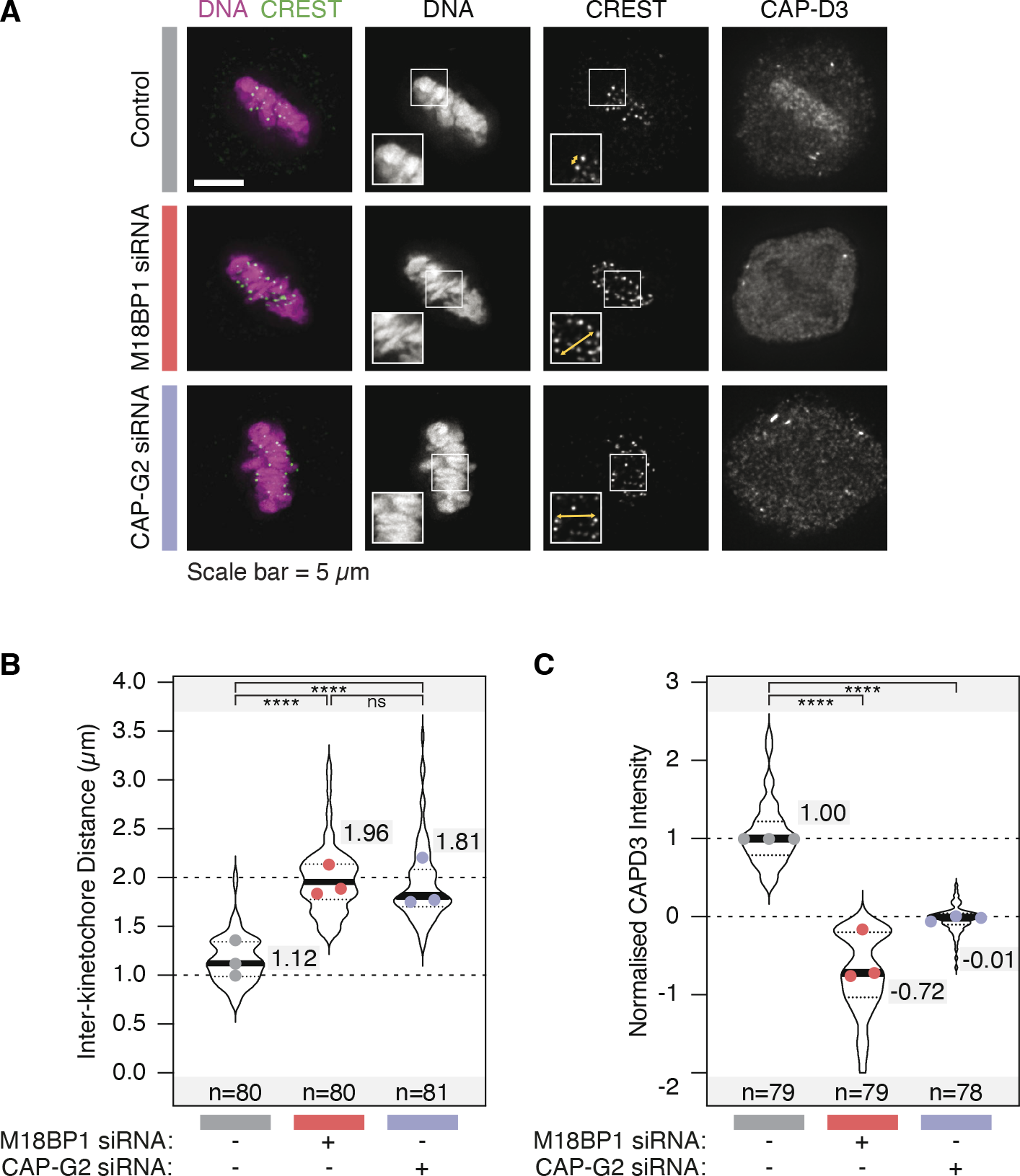
M18BP1 depletion during mitosis increases inter-kinetochore distances. (A) Example images of mitotic HeLa cells in metaphase treated with M18BP1 or CAPG2 siRNA. The yellow arrows in the insets highlight the increased inter-kinetochore distance upon depletion of either M18BP1 or CAPG2. (B) Quantification of inter-kinetochore distances from A. (C) Quantification of CAPD3 intensity from cells from A. “n” represents the number of cells analysed across three independent experimental repeats. The median of the combined data is shown for each condition and dots show the median of each experimental repeat.

**Figure S3.4:**
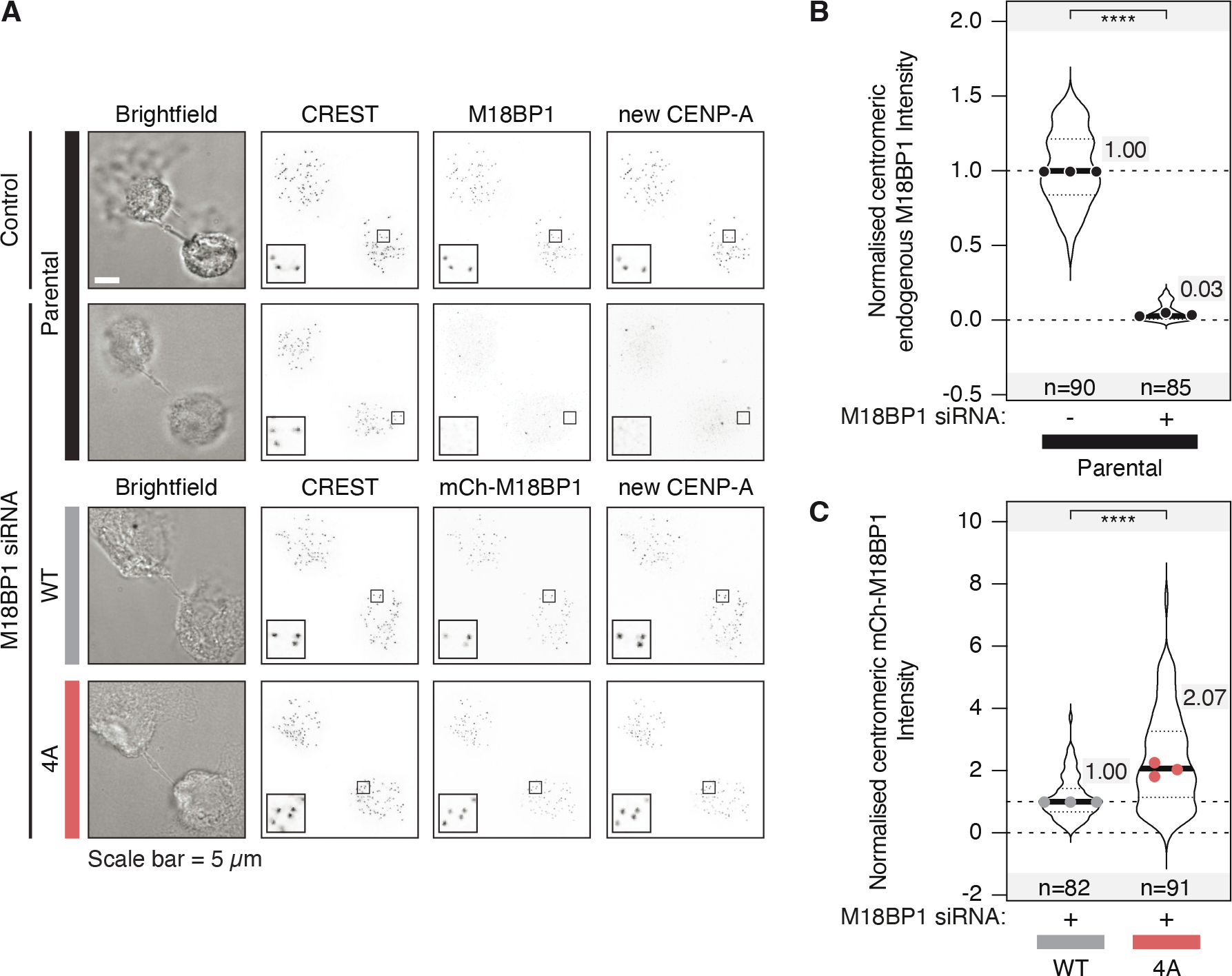
M18BP1 binding to condensin II is not important for new CENP-A deposition. (A) New CENP-A and mCherry-M18BP1 levels in early G1 cells expressing M18BP1 (WT or 4A mutant) upon depletion of the endogenous M18BP1. (B) Quantification of the endogenous M18BP1 levels in cells from A. (C) Quantification of mCherry-M18BP1 levels in cells from A. “n” represents the number of cells analysed across three independent experimental repeats. The median of the combined data is shown for each condition and dots show the median of each experimental repeat.

**Figure S4.1:**
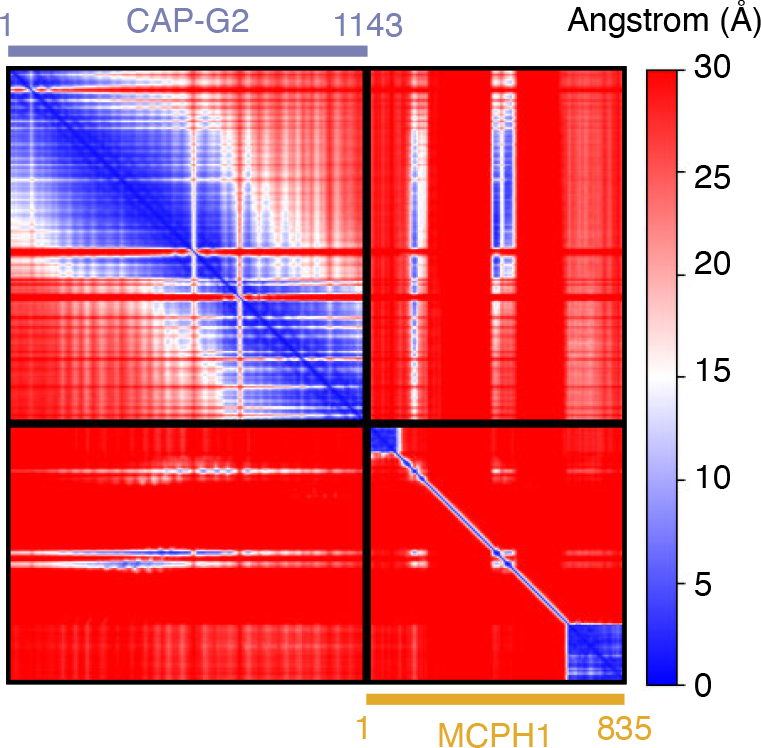
MCPH1 – CAP-G2 alphafold2 multimer prediction statistics. Predicted aligned error (PAE) plot for the CAP-G2 - MCPH1 multimer prediction. Higher confidence regions are coloured in blue.

**Figure S5.1:**
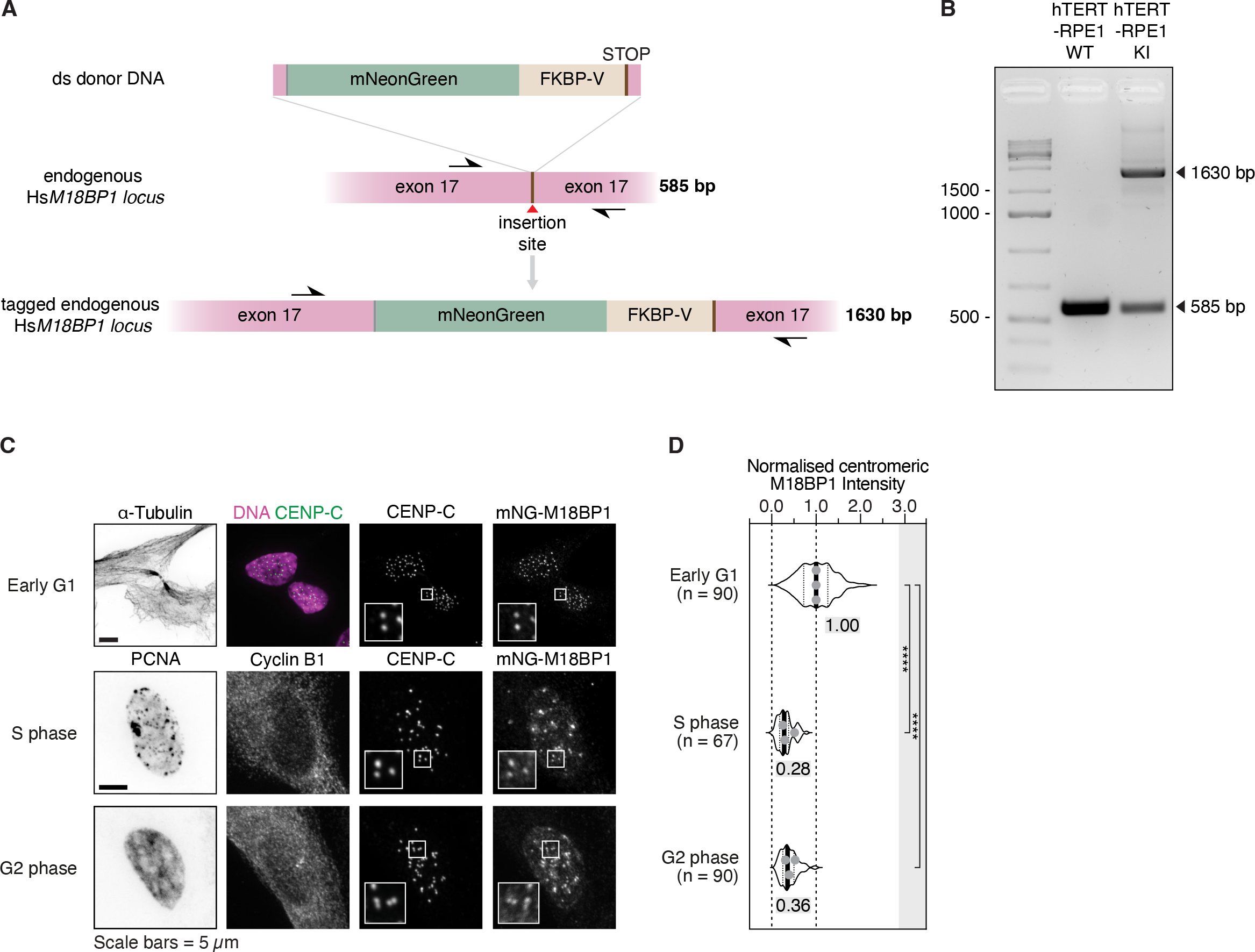
M18BP1 localises on chromatin and centromeres throughout the cell cycle. (A) Schematic illustrating the CRISPR/Cas9 knock-in strategy utilized to endogenously tag M18BP1 C-terminus (pink). Black arrows indicate the annealing sites of sequencing primers spanning the insertion site (red arrowhead). (B) Agarose gel electrophoresis (2% agarose) shows the integration of the donor DNA in the *locus* of interest. PCR amplicons represent untreated parental (lane #2) and M18BP1-mNG-FKBP-V heterozygous knock-in (KI) hTERT- RPE1 Flp-In TRex cell lines. (C) Localization of endogenously-tagged M18BP1 on chromatin and at centromeres during the cell cycle in RPE1 cells. Different cell cycle stages were identified using α-Tubulin, PCNA and CyclinB1 signals. (D) Quantification of centromeric M18BP1 levels from C. “n” represents the number of cells analysed across three independent experimental repeats. The median of the combined data is shown for each condition and dots show the median of each experimental repeat.

**Figure S5.2:**
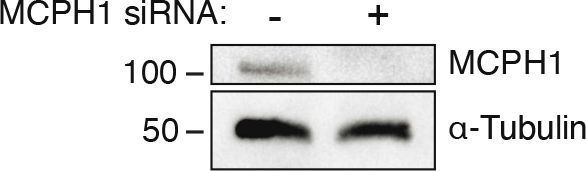
Depletion levels of MCPH1. Western blot shows the depletion levels of MCPH1 in HeLa cells upon siRNA treatment for 48h. The blot is representative of the depletion efficiency across experiments.

**Figure S5.3:**
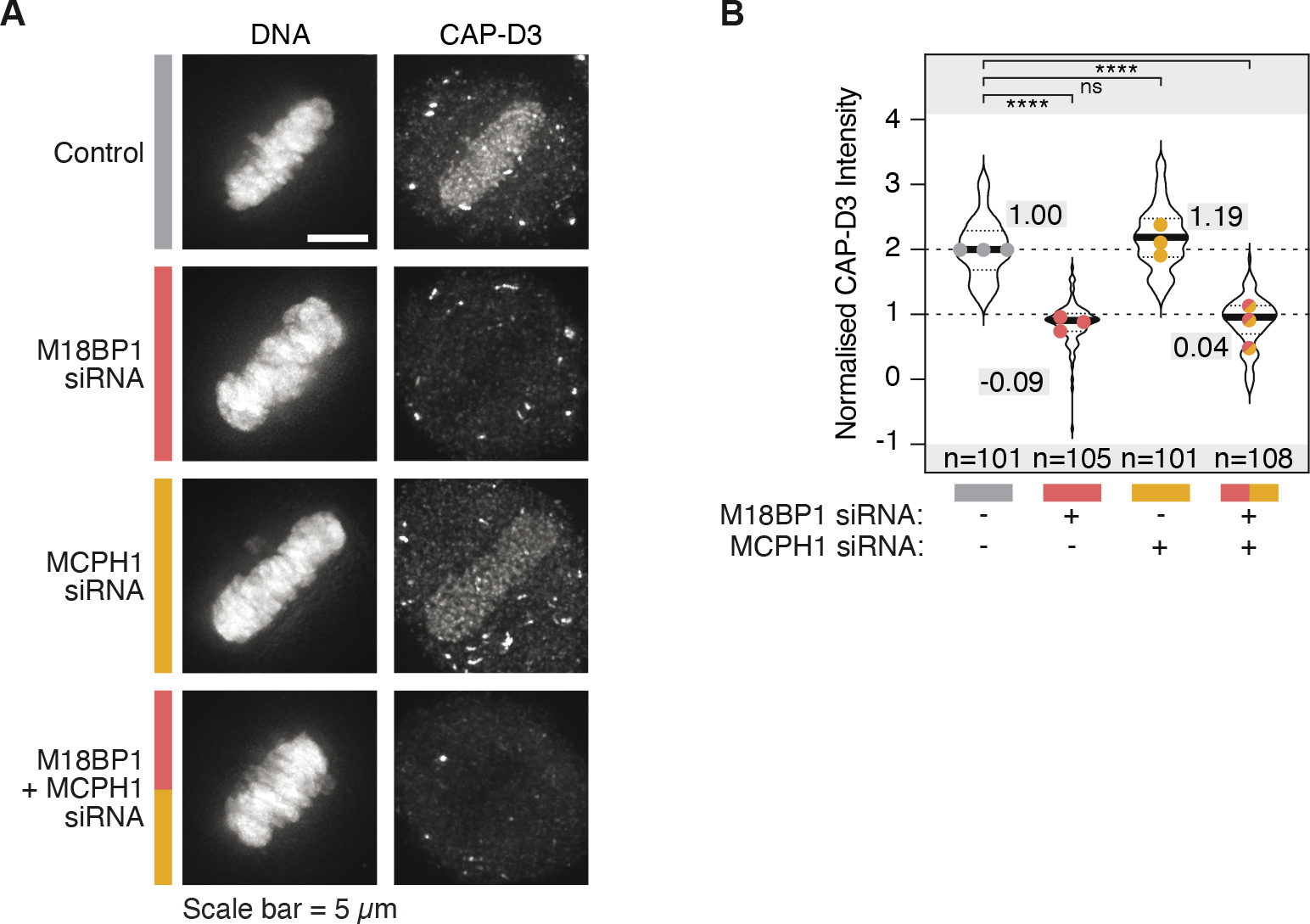
M18BP1 is essential for condensin II localization to chromatin during mitosis. (A) CAPD3 levels in mitotic HeLa cells in metaphase treated with the indicated siRNA. (B) Quantification of CAP-D3 signal from cells from A. “n” represents the number of cells analysed across three independent experimental repeats. The median of the combined data is shown for each condition and dots show the median of each experimental repeat.

**Figure S6.1:**
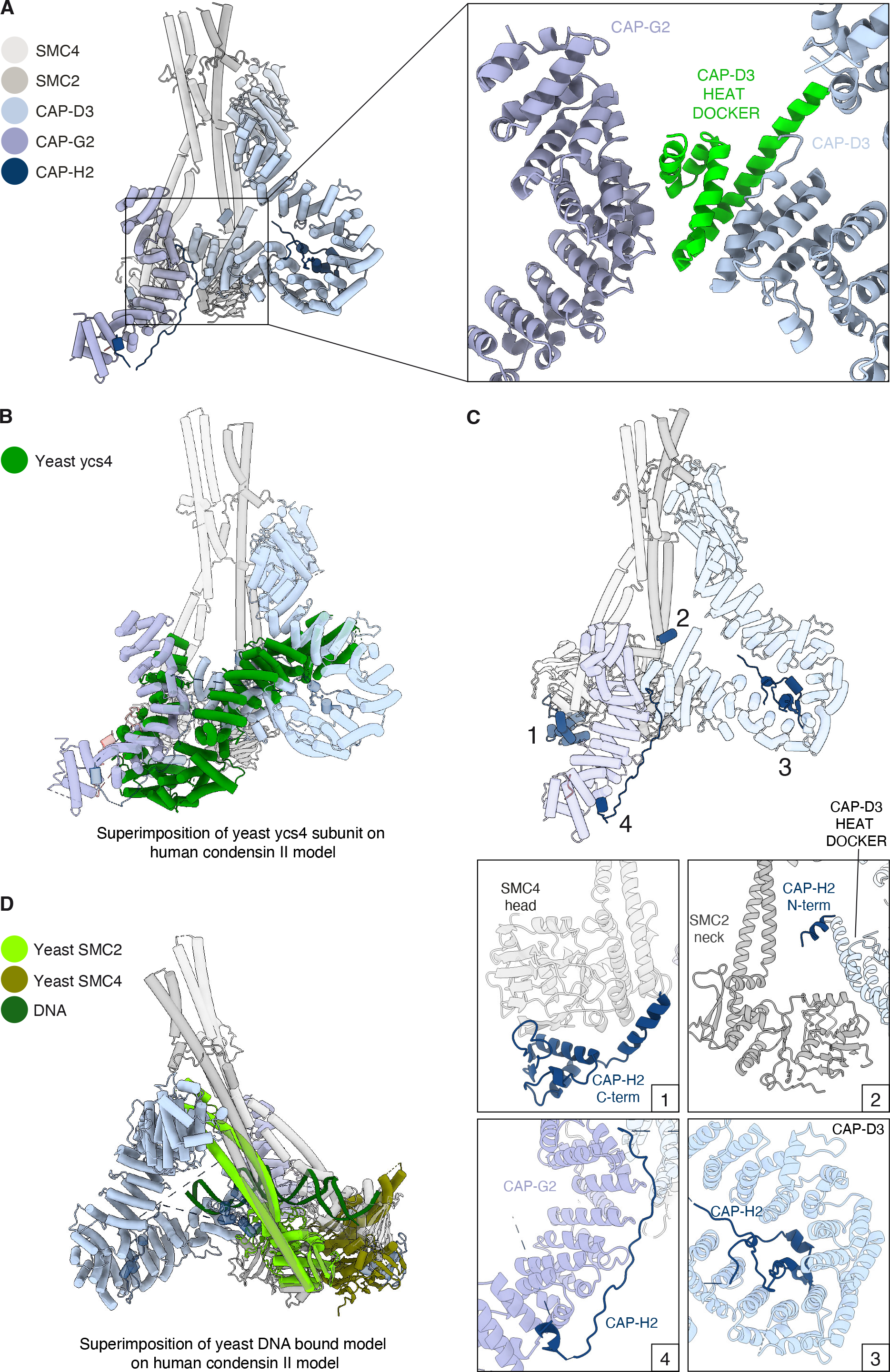
Condensin II holocomplex structure. (A) Cryo-EM model of apo Human condensin II holocomplex displacing SMC subunits in grey, CAP-D3 in light blue, CAP-G2 in purple and CAP-H2 in dark blue. Right panel: zoom on interaction between CAP-G2 and CAP-D3 subunits with “heat docker” residues (1178-1261) of CAP-D3 subunits coloured in green. (B) Superimposition of Yeast ycs4 coloured in green from apo not engaged yeast condensin (Lee B.G. *et al.* NSMB 2020, PDB code: 6YVU) with apo human condensin II. Superimposition was obtained aligning SMC2 subunits from both models. (C) Superimposition of apo Human condensin II with DNA bound yeast condensin (Lee B.G. *et al.* PNAS 2022, PDB code: 7Q2X). Model of apo human condensin II remains coloured as before, while subunits from Yeast condensin are coloured in shades of greens. Superimposition was obtained aligning SMC2 subunits from both models. (D) Model of apo human condensin II remains coloured as before with zoom ins on the interaction interfaces of CAP-H2 with (1) SMC4 head, (2) SMC2 neck, (3) CAP-D3 and (4) CAP-G2.

